# Three-dimensional markerless motion capture of multiple freely behaving monkeys for automated characterization of social behavior

**DOI:** 10.1101/2023.09.13.556332

**Authors:** Jumpei Matsumoto, Takaaki Kaneko, Kei Kimura, Salvador Blanco Negrete, Jia Guo, Naoko Suda-Hashimoto, Akihisa Kaneko, Mayumi Morimoto, Hiroshi Nishimaru, Tsuyoshi Setogawa, Yasuhiro Go, Tomohiro Shibata, Hisao Nishijo, Masahiko Takada, Ken-ichi Inoue

## Abstract

Given their high sociality and close evolutionary distance to humans, monkeys are an essential animal model for unraveling the biological mechanisms underlying human social behavior and elucidating the pathogenesis of diseases exhibiting abnormal social behavior. However, behavioral analysis of naturally behaving monkeys requires manual counting of various behaviors, which has been a bottleneck due to problems in throughput and objectivity. Here, we developed a three-dimensional markerless motion capture system that utilized multi-view data for robust tracking of individual monkeys and accurate reconstruction of the three-dimensional poses of multiple monkeys living in groups. Validation analysis in two monkey groups revealed that the system enabled the characterization of individual social dispositions and relationships through automated detection of various social events. Analyses of social looking facilitated the investigation of adaptive behaviors in a social group. These results suggest that this motion capture system will significantly enhance our ability to analyze primate social behavior.

## Main text

Non-human primates, including macaque monkeys, are social animals who utilize their knowledge about individual group members and their relationships to navigate a complex and dynamic social environment^1-6^. Together with their evolutionary proximity to humans, this makes monkeys a vital animal model for unraveling the biological mechanisms underlying human social behavior and elucidating the pathogenesis of neuropsychiatric disorders resulting in significant difficulties in social life^7,8^. However, to investigate the social functions and dysfunctions of non-verbal primates, it is necessary to assess their social interactions based on body actions, which are their primary communication tools^9,10^. Traditionally, behavioral actions are counted with visual inspection by human experts; however, the significant costs and reproducibility problems associated with manual annotation have been a major obstacle for these analyses. Recently, markerless motion capture using deep learning has been expected to overcome this issue as it allows high-throughput quantification of actions automatically and reproducibly^11^.

However, analysis of the social behavior of macaque monkeys in groups by utilizing a markerless motion capture has not been achieved. Since monkeys move freely in their living environment in a three-dimensional (3D) manner, 3D tracking and pose estimation of multiple monkeys are essential for detecting their social interactions. Previous reports on a primate 3D markerless motion capture system^12-14^ did not implement a multi-animal tracking algorithm and thus cannot be applied to groups of monkeys. Moreover, simple triangulation with single-view (2D) markerless motion capture^15,16^ is inappropriate because frequent and severe occlusions of monkeys hinder the tracking of individuals. Therefore, multi-animal tracking algorithms in a 3D space are required to reduce failures in individual tracking.

Here, we constructed a new pipeline utilizing multi-camera (multi-view) data for robust tracking and 3D pose reconstruction of multiple monkeys in their living environment. Applying this system to groups of Japanese macaques demonstrated that it could detect various social interaction events and extract their social characterization and relationships automatically. Furthermore, quantitative motion data enabled detailed analyses of social looking, a critical component of monkey social behavior that is difficult to analyze by visual inspection^1,9,17^.

## Results

### Construction of the pipeline for the 3D motion capture of multiple monkeys

We acquired multi-view videos of monkeys in their living environment from eight synchronized cameras set in a large (4 × 4 m) room of a semi-outdoor group cage (Fig S1). The monkeys wore a necklace-type color tag for individual identification (Fig S1b). Since the group cage had an adjoining small temperature-controlled room without cameras, the monkeys were often out of sight of any camera, thereby breaking the continuity of individual tracking (Fig S1a).

The overview of the pipeline integrating multi-view information for robust multi-animal tracking is summarized in Fig 1a. In the 2D video processing (Fig 1b), we trained and used three deep neural networks to detect the monkeys, estimate the 2D keypoint locations (e.g., nose, knee), and identify each detected monkey in each video frame. In addition, we used a box tracking algorithm to track the monkey detections across video frames, resulting in fragments of continuous monkey tracks in the video from one camera (single-view tracklets). To integrate multi-view information, the monkey detections in different views associated with the same individuals were searched (cross-view matching) based on the geometric (keypoint) and appearance affinities of pairs of detection^18^ (Fig 1c). Since the appearance of the monkeys used in this study was very similar, we used the ID (color tag) detections in corresponding single-view tracklets to calculate the appearance affinities between the monkey detections. To reduce computational load, we performed cross-view matching for one keyframe every 0.5 s. The connections between cross-view matched monkey instances across time were estimated by maximizing the consistency of single-view tracklets (Fig 1d). This process resulted in multi-view tracklets, i.e., sets of 2D monkey detections corresponding to the same monkeys across views and video frames (colored lines in Fig 1d). Then, the IDs were assigned to each multi-view tracklet based on the results of ID detection in 2D video processing. Finally, we reconstructed the 3D motion of each identified monkey from the ID-assigned multi-view tracklets using Anipose^19^. A representative example of 3D motion capture is shown in Fig. 1e and Movie S1.

**Figure 1:**
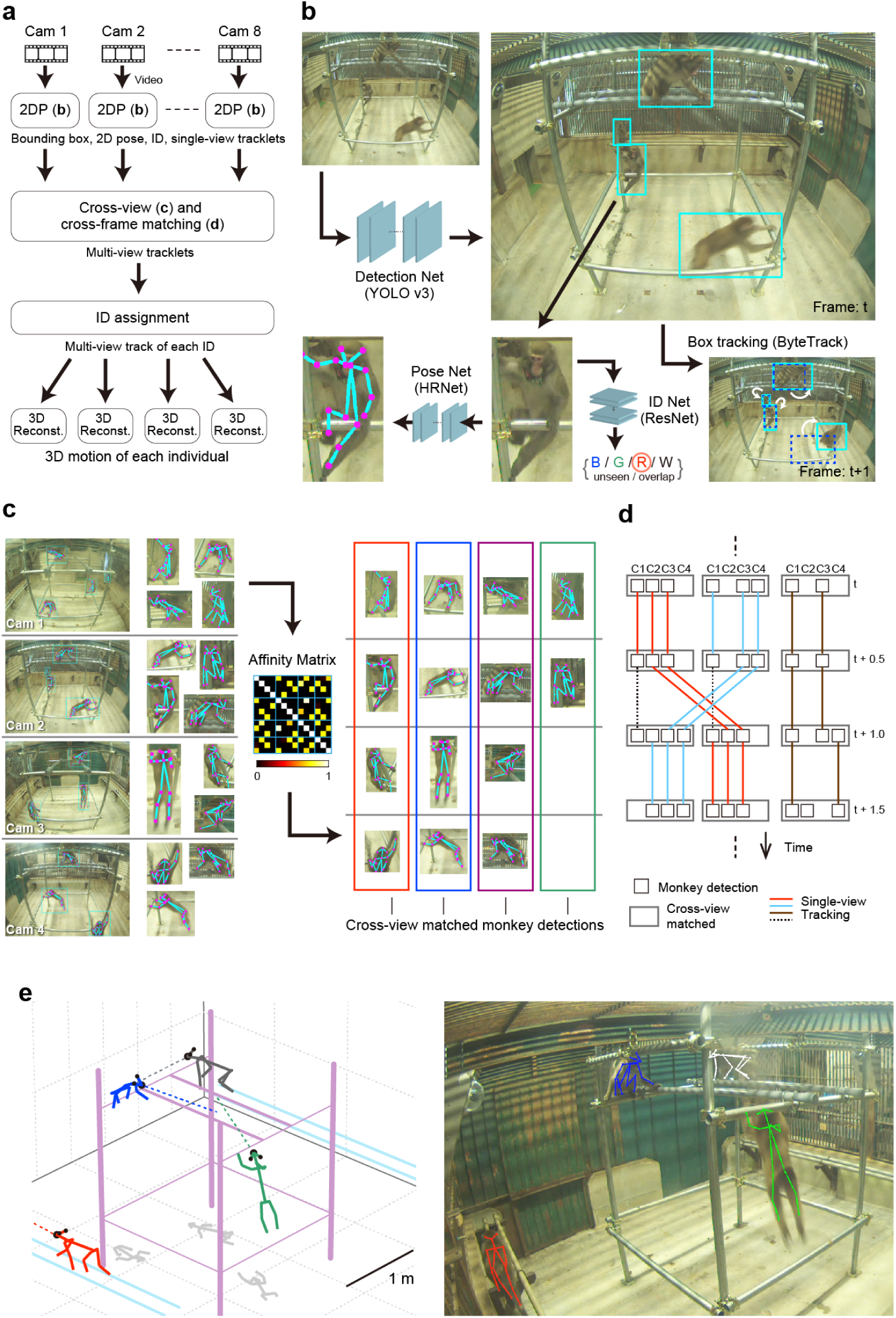
Motion capture algorithm. (**a**) Overview of the data processing pipeline. (**b**) 2D processing of a video from a camera. Three deep neural networks (Detection Net, Pose Net, ID Net) and a box tracking algorithm were applied. Parentheses indicate the names of the algorithms. ID Net output either monkey ID (color of the tag, blue [B], green [G], red [R], white [W]), *unseen* (the color tag was unseen), or *overlapped* (multiple monkeys overlapped in the image). (**c**) Cross-view matching. Left: example monkey detections in each view (images from four cameras are shown for simplicity; left side, whole images; right side, zoomed images of the detections). Center: affinity matrix representing the affinities of all pairs of detections (14 × 14 detections with the example shown in the left panel). Optimal matching was found based on the affinity matrix^18^ (right). (**d**) Cross-frame matching. C1-4, cameras 1-4. The cross-view matched instances (wide rectangles) were connected across frames based on single-view tracking (solid lines). The dotted lines indicate the single-view tracks that were excluded because of inconsistencies. (**e**) An example of a 3D motion capture result. Left: 3D plot. Black dots indicate ears and noses. Dotted lines indicate facial direction. Line colors indicate the IDs of the monkeys. Gray lines on the floor are shadows (horizontal coordinates) of the keypoints. Right: 3D posture reprojected on the corresponding image.

### Performance validation in monkey groups

We recorded and analyzed the social behaviors of two groups of monkeys (Table S1). Each group included four monkeys wearing a blue (B), green (G), red (R), or white (W) ID tag. To evaluate the accuracy of the constructed system, we computed the errors across 3D poses estimated by the markerless motion capture and those obtained by the annotation of a human experimenter. We assessed the errors for the keypoints and face direction, which was calculated from the locations of the nose and ears (Fig 2a). All median errors of the keypoint locations were <50 mm, while that of face direction was 14.0°, indicating good performance in pose estimation. We also checked the system’s performance for individual tracking by comparing the estimated IDs to those annotated by a human experimenter. The ID precision (IDP; correctness of estimated IDs) and recall (IDR; accuracy in recovering IDs) were 97.5 ± 1.3% and 86.0 ± 2.8% (mean ± s.e.m.), respectively, suggesting high performance. The slightly smaller IDR may be due to short tracklets without ID detection, e.g., shuttling between the recording and temperature-controlled (non-recording) rooms.

**Figure 2:**
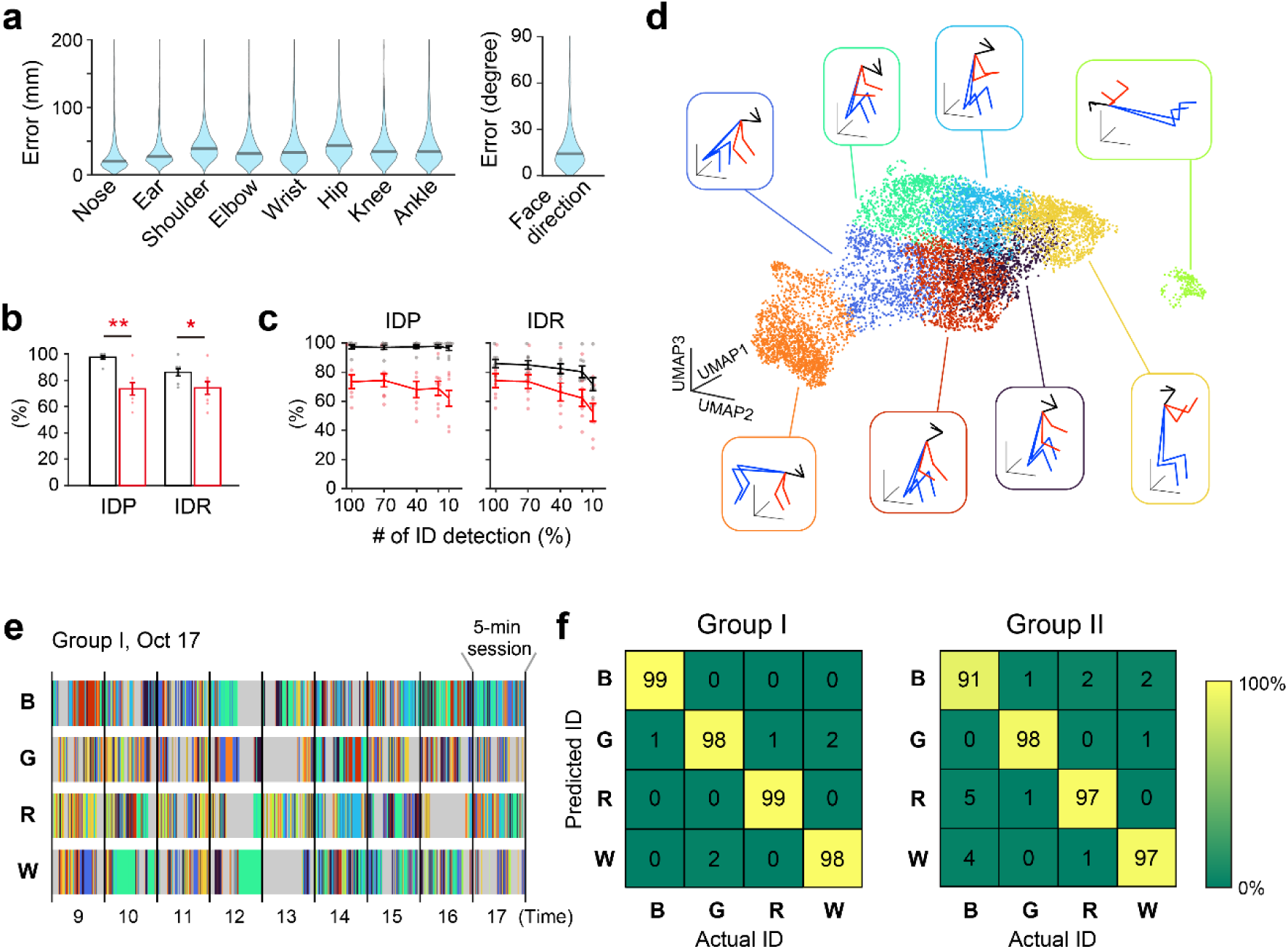
Performance validation. (**a**) Distribution of errors across 3D postures estimated by the markerless motion capture system and those by human annotation. Black line, median errors. *n* = 548 monkeys. Errors of the corresponding left and right body parts were averaged. (**b**) Mean ID tracking precision (IDP) and recall (IDR) of the proposed (black bars) and control (red bars) algorithms. Error bars indicate standard error of the mean (s.e.m.). *n* = 8 5-min recordings. \**p* < 0.05, \*\**p* < 0.01, paired *t*-test. (**c**) Mean IDP and IDR after random reduction in the frequency of ID detection. ID detection in each view was reduced by randomly masking 1-s time bins. Black and red lines indicate the performance of the proposed and control algorithms, respectively. Error bars indicate s.e.m. *n* = 8 5-min recordings. (**d**) Uniform Manifold Approximation and Projection (UMAP) projection of monkey postures. The colors of the dots indicate the result of classification using the *k*-means algorithm (*k* = 8). The inset indicates the mean posture of each cluster. (**e**) The example posture patterns of four monkeys (blue [B], green [G], red [R], and white [W]) in Group I on 1 day. A 5-min session is shown every 1 h. Colors correspond to the classes in d. Gray indicates that the monkey was not detected. (**f**) The confusion matrix of support vector machine performance predicting a monkey ID based on its posture pattern.

To assess the effectiveness of multi-view information in tracking, we compared its performance with a conventional approach that does not utilize multi-view information but reconstructs the 3D motion of a monkey tracked separately in each view (control algorithm). The IDP and IDR of the control algorithm dropped by approximately 20% and 10%, respectively, compared to the proposed algorithm (Fig 2b); its poor performance was partly due to erroneous integration of different monkey detections across views because of ID tracking errors (ID switch) in some views. We also tested tracking performance after randomly reducing the frequency of ID detection to check the system’s potential for applications in which ID detection will be sparser, e.g., field recordings where severe occlusions are common or use of faces for ID instead of color tags. We found that the performance of the proposed algorithm was maintained well compared with the control algorithm (significant interaction [IDP, *p* = 0.038; IDR, *p* = 0.041] between the algorithms and the ID detection rate in two-way repeated measures ANOVA), suggesting our system was robust to reductions of the ID detection rate, thanks to the utilization of multi-view information for tracking (Fig 2c).

To verify the practicality of this motion capture system, we attempted to identify individuals by their posture patterns. The postures were classified in an unsupervised manner using *k*-means clustering after dimension reduction (Fig 2d, e), and the frequency and duration of each posture were used to discriminate individuals by a support vector machine (SVM). We found that the SVM could discriminate individuals with an accuracy of >95% (98.5 ± 0.3% and 95.8 ± 1.6% in Groups I and II, respectively; Fig 2f), suggesting that our system is sufficiently accurate to extract each individual’s movement characteristics.

### Automatic detection of social behavioral events and behavioral characterization

Then, we analyzed the social behaviors of two groups of monkeys (Groups I and II; Table S1). The analyses were focused on the last 8 recording days in the 3 (Group I) and 4 weeks (Group II) stay in the cage. In this study, we defined and automatically counted affiliative (*Proximity*, *Groom*), vigorous (*Chase*, *Glare*, *Grab*, *Pounce*), and other (*Mount*, *Look*) social behaviors based on quantitative motion parameters, e.g., distance between two monkeys and their postures (Fig 3a; see also Methods and Text S1 for detailed definitions). Note that we defined the Look as orienting the head toward a conspecific according to previous studies^3,15,20^ since measuring the actual gaze direction based on the eye movements of freely moving monkeys is difficult in our recording setup. Comparisons between the automatic event detections and manual event annotations by human experts for Chase (recall: 70%; precision: 76%; *n* = 23 events detected by the experts), Groom (recall: 74%; precision: 83%; *n* = 68 events), and Mount (precision: 91%; *n* = 10 events; recall could not be calculated because Mount was an infrequent event) indicated high detection performance for these events (see Movie S2 for examples of the events).

**Figure 3:**
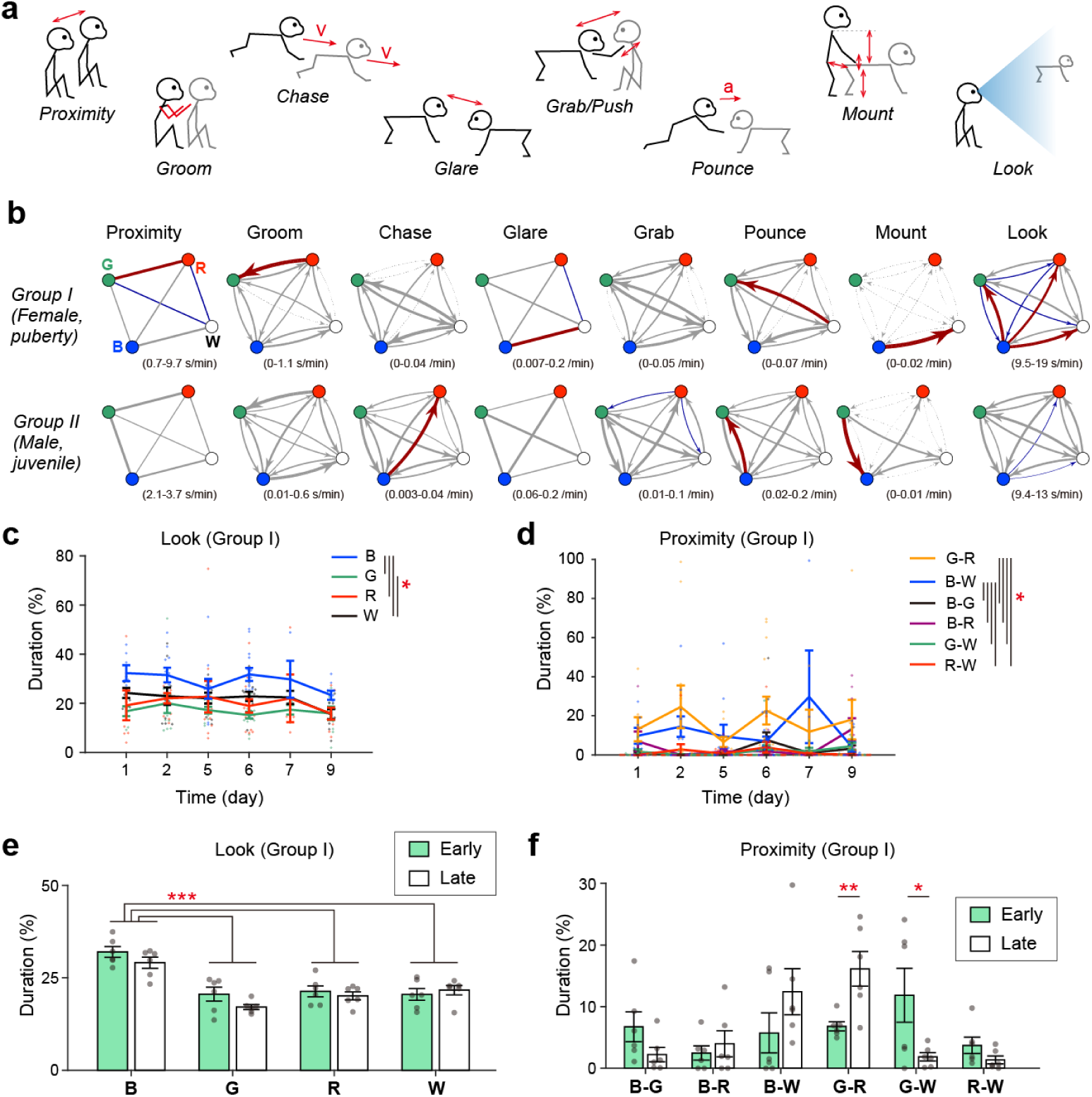
Analysis of social behavioral events. (**a**) Illustrations of definitions of social behavioral events detected automatically from the motion data. The double arrow indicates the distance between the pair. *v*, velocity. *a*, acceleration. Detailed definitions are described in Text S1. (**b**) Average frequencies and durations of social behaviors between each pair of monkeys. The data are summarized as an undirected or directed graph, where each node represents an individual, and the edge thickness represents the averaged frequency or duration of the event. Red and blue indicate that the value was significantly larger and smaller than the individual shuffled data, respectively (*p* < 0.05 with Bonferroni’s correction). Parentheses indicate the ranges of values in the graphs. *n* = 58 and 111 sessions in Groups I and II, respectively. (**c, d**) The mean duration of Look (c) and Proximity (d) in each analysis day in Group I. Days 3, 4, and 8 were excluded from the analysis since the number of available samples (recording sessions) was low (0, 1, and 2, respectively; Table S2). \**p* < 0.05, Bonferroni’s method. *n* = 7, 11, 10, 14, 4, and 9 sessions on days 1, 2, 5, 6, 7, and 9, respectively. (**e**) Comparison of the mean duration of Look between the early and late phases of the stay in the recording cage in Group I. \*\*\**p* < 0.001, Bonferroni’s method. *n* = 6 days in each phase. (**f**) A similar comparison on the mean duration of Proximity. \**p* < 0.05, \*\**p* < 0.01, simple main effect analysis. Error bars in c–f indicate standard error of the mean. See Table S3 for the corresponding analysis of variance results of c–f.

Automated counting of each event revealed the unique characteristics in the social disposition of individuals and groups (Fig 3b). In the female puberty group (Group I), Monkey B, the smallest in the group, tended to orient her head to the other individuals more frequently. On the other hand, Monkey R, which originated from a different colony to the others, tended not to participate in vigorous behavior but showed affiliative behavior to Monkey G. In the juvenile male group (Group II), the counts of vigorous behaviors were high, consistent with the visual inspections that playing as a whole was common in this group. Pair- or individual-specific social behaviors were fewer in Group II than in Group I, but were still detected.

To examine the stability of the detected behavioral characteristics, we calculated the mean duration of Look and Proximity for each analysis day in Group I, in which significant individual differences of those events were found (Fig 3c, d). The results showed stability throughout the 8 successive recording days (significant main effect of ID or pair, but no significant main effect or interaction relating to the recording date in two-way repeated measures ANOVA; see Table S3 for the ANOVA results), suggesting the analysis could extract the social characteristics of each individual or pair. In addition, the tendency of Monkey B to perform looking behavior frequently was preserved across the initial 7 recording days (early phase) and the last 8 recording days (late phase) (Fig. 3e). Conversely, the proximity duration of the G-R and G-W pairs was significantly increased (*p* = 0.0094) and decreased (*p* = 0.048), respectively, in the late phase, suggesting a proximity pair transition (Fig 3f). These results indicate that the system may be helpful in tracking long-term changes in the social relationships within monkey groups.

We then examined whether individuals could be discriminated from the patterns of their social behavior by SVM (Fig 4a, b) in the same way as with the posture patterns (Fig 2f). The results indicated high prediction accuracy (91.8 ± 5.7% and 80.0 ± 6.4% in Groups I and II, respectively), although accuracy was slightly lower than with posture patterns, especially in the male juvenile group (Group II). To examine which social behavioral events were important for individual discrimination, we evaluated the performance of the SVM model when only a single type of event was used and when a single type of event was missing. The performance of the SVM models using a single type of event (Fig 4c) indicated that many different events could individually predict the monkey ID above chance level, although performance was lower than with the full SVM model using all events. On the other hand, missing a single type of event did not result in a drop in discrimination performance, except for Look (Fig 4d). These results indicate that the social behavioral events analyzed here, especially Look, were effective in characterizing individual monkeys. Furthermore, ignoring the ID of the social behavior partner decreased discrimination performance (Fig 4e), indicating that social relationships were important for ID discrimination. The overall results suggest that our system has the potential to detect the social behavioral characteristics of individuals and their relationships.

**Figure 4:**
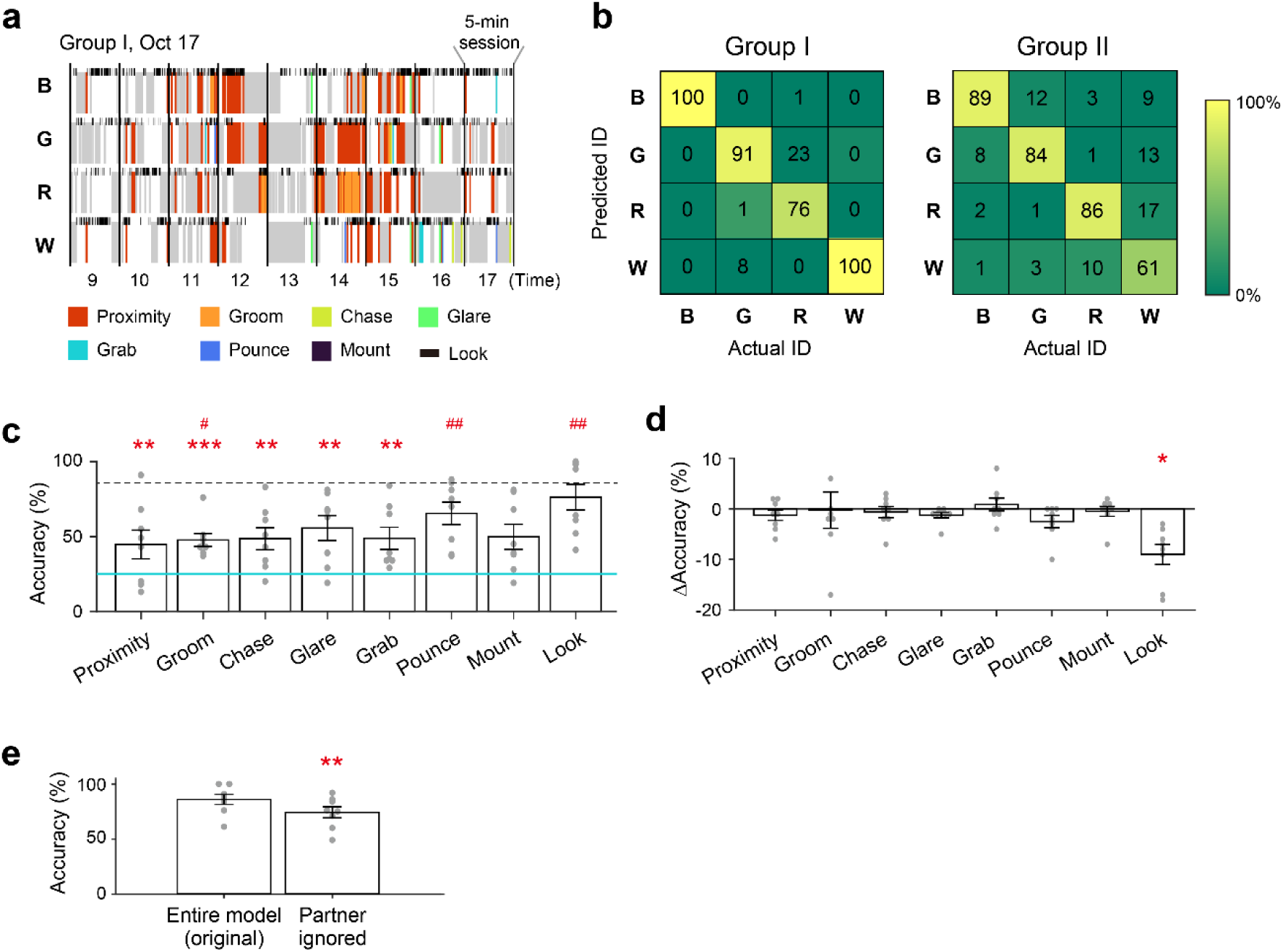
Individual discrimination based on the pattern of social behavior. (**a**) The social behavioral patterns of monkeys in a group on 1 day. The same period as Fig 2e is shown. Colors correspond to social events. Gray indicates that the monkey was not detected. White indicates that none of the events were detected. Timings of Look events are shown on the top of each row with narrower markers. Note that the partners of each social behavior are not presented in this graph, although partner information was used in b–d. (**b**) The confusion matrix of support vector machine (SVM) performance predicting a monkey ID based on its social behavioral pattern. (**c**) Mean prediction accuracy of SVM models using only a single type of social behavior. The dotted line indicates the mean accuracy of the entire (original) model (b). \**p* < 0.05, \*\**p* < 0.01, \*\*\**p* < 0.001, significant difference from the entire model using a paired *t*-test with Bonferroni’s correction. #*p* < 0.05, ##*p* < 0.01, ###*p* < 0.001, significant difference from the chance level (25%, solid blue line) using a one-sample *t*-test with Bonferroni’s correction. (**d**) The mean difference in the prediction accuracy of SVM models missing a single type of social behavioral event from those of the entire model. \*\**p* < 0.01, significant difference from the entire model using a paired *t*-test with Bonferroni’s correction. (**e**) Comparison of the mean accuracy between the entire model and the model ignoring social behavior partners. \**p* < 0.05, paired *t*-test. Error bars in c–e indicate standard error of the mean.

### Analysis of social looking

Looking at a conspecific is a critical component of monkey social behavior^1,9^. We further analyzed Look behavior, which was the most effective behavioral event for the individual discrimination above. First, we compared the Look duration calculated with and without shuffling the monkey motion data across recording sessions (Fig S2a). The actual duration of Look behavior was significantly larger than that with the shuffled data, supporting the tendency to look at other monkeys. We also calculated Look duration with temporally shifting monkey motion data (Fig S2b) and found that the peak was at zero time shift, suggesting that looking follows the target monkey’s movement. Counting the third party’s Look behavior toward each social behavioral event demonstrated that the monkeys tended to observe vigorous behavior, such as Chase (Fig 5a), and a detailed analysis of Chase demonstrated that the third party’s facial direction followed the chasing movement (Fig 5b). Such social looking is essential for understanding ongoing situations and relationships between group members^1-5,21^.

**Figure 5:**
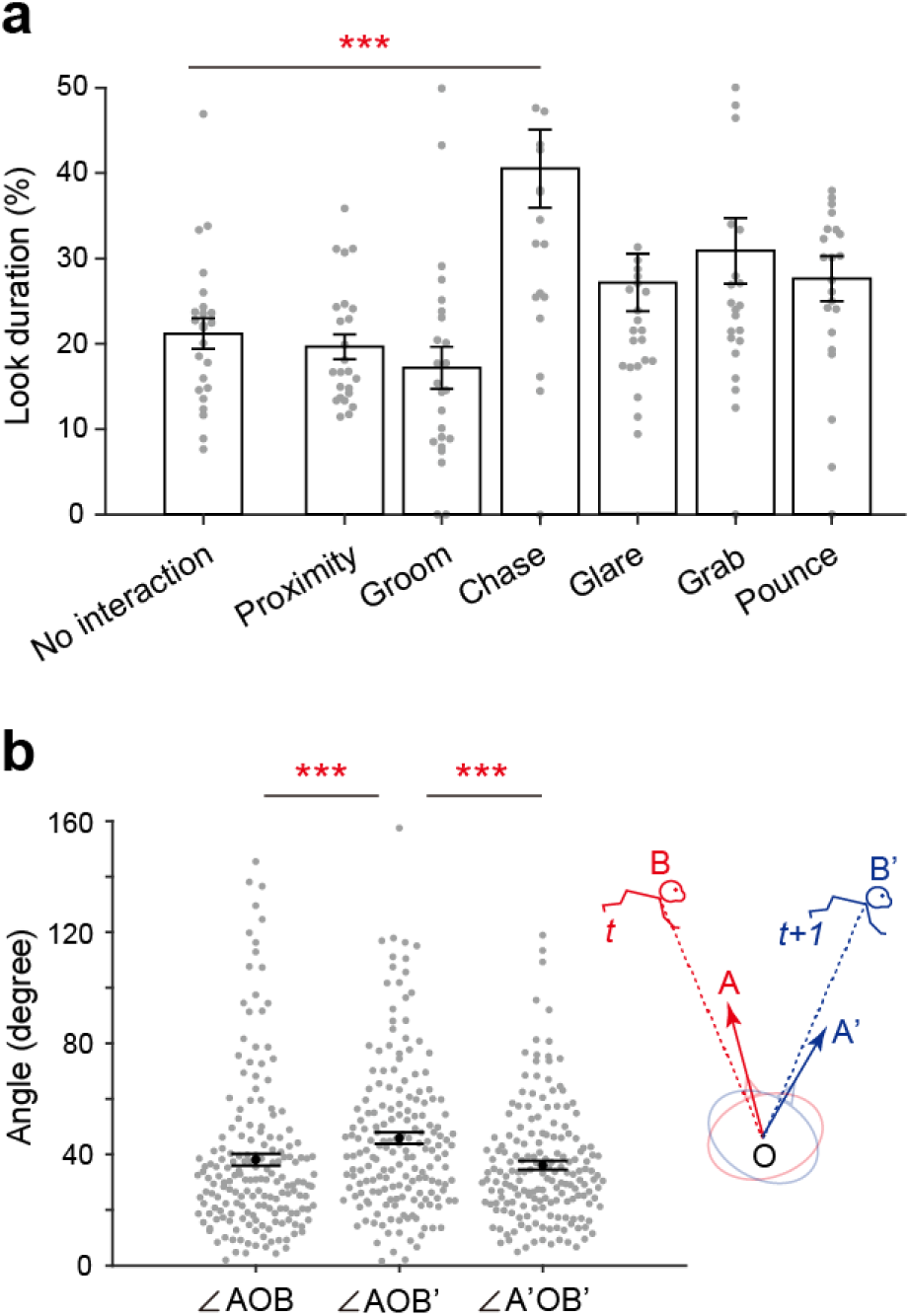
Third-party observation of social behavior. (**a**) Mean Look duration in 1-s time bins containing each social behavioral event. No interaction, the time bin contained no social behavioral event detection. \*\**p* < 0.01, paired *t*-test with Bonferroni’s correction comparing No interaction and another event. *n* = 24 pairs of monkeys. (**b**) Distribution of angles between the third-party’s face direction and the direction of the target monkey during Chase. The inset shows an illustration of the parameters. A and A’, face directions at the beginning and end of a 1-s bin, respectively. B and B’, the target monkey location relative to the third-party monkey location (O) at the beginning and end of a 1-s bin, respectively. *n* = 178 1-s bins including Chase and third party’s Look. \*\*\**p* < 0.001, paired *t*-test with Bonferroni’s correction. Black dots, mean angle. Error bars in a and b, standard error of the mean.

In primates, mutual looking (both parties look at each other) has social meaning, as direct staring is a threatening behavior, and gaze aversion is often associated with anxiety and submissiveness^9,22^. We compared the actual mutual look duration with the chance-level duration that would be expected if each monkey’s looking behavior was independent (Fig S2c). The actual mutual look duration was significantly shorter than the chance level, suggesting monkeys may avoid mutual looking. To analyze which monkey showed stayed or withdrawn looking at the end of the mutual look event, we counted the stay ratio among all mutual look events for each pair of monkeys (Fig 6a, b). The result revealed that one monkey often withdrew from looking at most of the members in each group (Monkeys B and W in Groups I and II, respectively), one monkey stayed in looking at all members in Group II (Monkey B), and two monkeys withdrew or stayed depending on the member in Group II (Monkeys G and R). Interestingly, when we analyzed the relationship between the stay rate and total Look duration in pairs in each group (Fig 6c), negative correlations were found in both groups. Previous studies reported that while subordinate monkeys tend to avert their gaze from dominant monkeys when two monkeys face each other^22^, the dominance rank is negatively correlated with looking duration^17^. Thus, the stay-withdraw pattern we found might reflect the monkeys’ hierarchy. These results suggest the utility of analyzing looking behavior based on the motion capture system for investigating monkeys’ adaptive behaviors in a social group.

**Figure 6:**
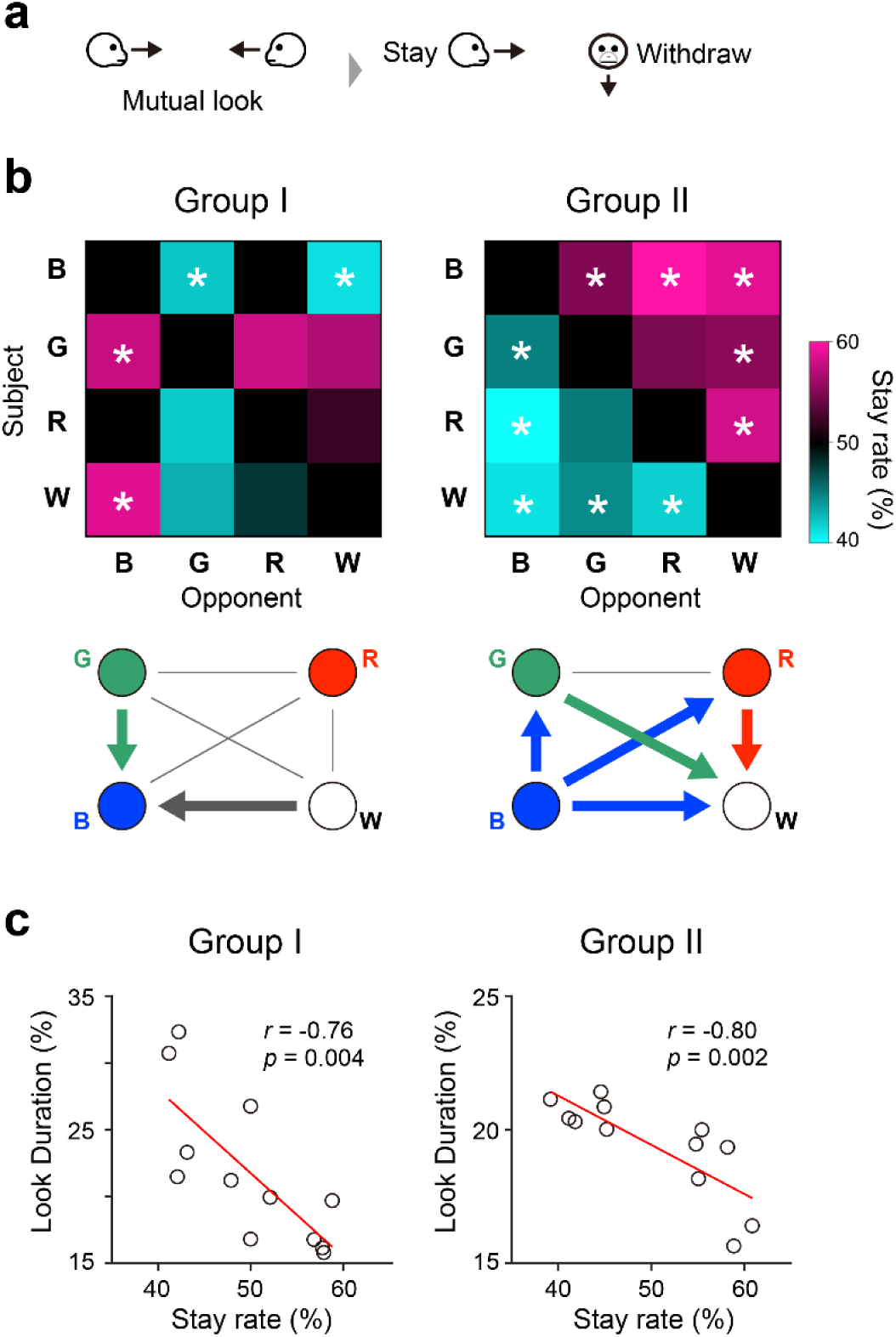
Face orientation response to mutual looking may reflect hierarchy. (**a**) Illustration of the face orientation responses. After each mutual look bout of a pair, we counted which monkey stayed looking (Stay) or withdrew (Withdraw). (**b**) Top, the number of Stay events among all mutual look events for each pair of monkeys. \**p* < 0.05, binomial test compared with the chance level (50%) with Bonferroni’s corrections. Bottom, schematics summarizing the significant Stay-Withdraw relationships. An individual at the tail of an arrow (also corresponding with the arrow’s color) has a significantly larger Stay rate than chance in the bout with the opponent at the head of the arrow. (**c**) Relationship between the Stay rate and total Look duration in monkey pairs. *n* = 12 permutations of pairs per group. *r* and *p*, Pearson’s correlation coefficient and corresponding *p*-value.

## Discussion

Here, we constructed a pipeline for the long-term 3D markerless motion capture of monkeys living in groups. The pipeline utilized multi-camera (multi-view) data for robust tracking of individual monkeys and accurate reconstruction of their 3D poses. Using this system, we obtained 3D motion data of monkeys living in groups and analyzed social behavior based on motion data for the first time. Our analysis demonstrated that this system could characterize individual motion and social traits and define their relationships. We further demonstrated the system’s usefulness for analyzing the adaptive behaviors of monkeys in social groups through the detailed analysis of their looking behavior.

Measuring monkeys’ 3D poses and motion in groups enabled the analysis of their various social interactions. In contrast to image classifications using supervised machine learning for detecting a specific behavior with large amounts of manually annotated training data^23,24^, 3D motion data can be used flexibly to detect various behaviors, including those that are difficult to detect with image-based analysis, e.g., looking. Previous studies also proposed 3D markerless motion capture systems for monkeys^12-14^ and demonstrated detailed and automatic analysis of various behaviors of freely moving monkeys. However, they cannot be applied to monkeys in a group due to the lack of a multi-animal tracking algorithm. Our system overcomes this limitation through robust individual tracking using multi-view data (Fig 1c, d).

Our system used an optimization algorithm for cross-view matching, but machine learning algorithms for cross-view matching have also been suggested^25^. An important advantage of the optimization-based approach is its ease of extension. The development of 2D processing deep learning algorithms is more active than for 3D algorithms (paperswithcode.com) because of their high versatility. In addition, many existing training datasets^26-28^ were made for 2D image processing. It would be relatively easy for our system to be applied to existing datasets of different species. Moreover, updating the 2D processing algorithm or ID detection algorithm would directly enhance the performance of our pipeline.

Social looking is a critical component of monkeys’ social behavior and has received much attention in studies on primate social behavior. Monkeys understand the relationship between others through observations^1-3,5,21^. Their visual attention toward others depends on their social relationship^17,22^, and gaze and gaze aversion may be social signals by themselves^9,22^. Neuroscience studies in laboratory settings have revealed the neural mechanisms involved in monkeys’ looking behavior^6,7,29^ and their impairment in animal models of autism^30,31^. Furthermore, the neural bases of sophisticated social functions in monkeys have been mainly studied in highly controlled laboratory settings while monkeys’ movement is constrained^6^. Although these studies have provided many important insights into the neural basis of social behaviors and their dysfunctions^6,7,29,32^, such approaches have problems of external validity. Examination in a more naturalistic (ethologically relevant) social environment is needed to compensate for this limitation^33-39^. Detailed quantitative analysis of social behavior, including social looking, of freely behaving monkeys in groups with markerless 3D motion capture will provide unique opportunities to extend findings in a specific social task in the laboratory.

In addition, markerless 3D motion capture could be applied to analyze various social functions of primates, e.g., the long-term changes of relationships in groups, the developmental trajectory of social skills, and relationships in larger groups^10,38,40^. Our approach may also be applicable to field studies, which require the tracking of individuals with facial identification^38^, thanks to its robust tracking ability using multi-view data. Extensions for detecting other social gestures in different modalities, such as vocalizations and facial expressions^2,9^, and computational analysis of primate social behavior using motion data with high spatiotemporal resolution^41-43^ will be critical next steps.

## Methods

### Animals

Two groups (Groups I and II) of Japanese macaques (*Macaca fuscata*) were used in this study. Each group consisted of four monkeys of the same sex and similar age (Table S1). The experiments were approved by the Animal Welfare and Animal Care Committee of the Center for the Evolutionary Origins of Human Behavior of Kyoto University (2022-101) and conducted in accordance with the Guidelines for the Care and Use of Animals of the Center for the Evolutionary Origins of Human Behavior, Kyoto University.

### Video recording

A group cage consisting of one semi-outdoor large room (4 × 4 × 2 [height] m) and one small temperature-controlled room (2 × 1.5 × 2 [height] m) was used for recording (Fig S1). A small rectangular hole (0.7 × 0.7 [height] m) with an electric door connected both rooms, making it easy for the experimenters to clean each room by keeping the monkeys in the other room. We placed eight cameras (acA2040-35gc, Basler) equipped with a wide lens (ML410 4–10 mm, Theia) on the wall near the ceiling inside the large room surrounding the center of the room. Each camera was mounted in a custom stainless-steel housing with an acrylic dome window (O’Hara) securely fastened to the cage frame to prevent the monkeys from moving or touching the camera. Videos (2,048 × 1,536 px, 24 fps) were captured synchronously from the eight cameras using the Motif acquisition system (Loopbio).

### Camera calibration

Intrinsic (e.g., lens distortion coefficients) and extrinsic (camera pose and location) camera parameters are required to reconstruct a 3D coordinate of a keypoint (e.g., nose, shoulder, elbow) from 2D coordinates of the keypoint projected onto camera images from different views. To calibrate these parameters, first, we initialized the intrinsic parameters of each camera with the cv2.omnidir.calibrate() function in OpenCV^44^ using images of checkerboards from multiple angles. Then, the extrinsic parameters of each camera were initialized with the cv2.solvePnP() function from the known 3D coordinates in the cage, e.g., corners of the room, and their 2D coordinates projected onto the camera image. Finally, we waved a wand with a marker (ping-pong ball) on the tip throughout the recording volume, and the intrinsic and extrinsic parameters of all cameras were simultaneously optimized with a bundle adjustment^19^ by minimizing the reprojection errors of the marker locations.

### Data collection

Groups I and II were moved to the recording cage and recorded for 3 weeks in October and 4 weeks across May and June, respectively. Before moving to the recording cage, the monkeys had lived with the same group members for >10 months, either in a similar-sized group cage (Group I) or in a large breeding colony (Group II). The monkeys wore a colored necklace for individual identification (Fig S1b). The light in the large room was turned on from 08:30 to 17:30. Food pellets were supplied once a day in the food container (Fig S1a). Supplemental fresh vegetables and fruits were given 2–3 times a week. Water was supplied *ad libitum* from water dispensers (Fig S1a). Recording was conducted daily during the daytime from 08:00 to 18:00. Sometimes, recording was paused due to technical issues. Unless noted, all data used were from each group’s last 8 recording days after the monkeys were well acclimated to the recording environment.

### Two-dimensional video processing

Different deep neural network models were used for monkey detection (YOLOv3^45^), pose estimation (HRNet w32^46^), and monkey identification (ResNet50^47^) (Fig 1b). The monkey detection network (Detection Net) estimated bounding boxes around the monkeys in an image. The pose estimation network (Pose Net) estimated 15 keypoints (nose, left and right ears, shoulders, elbows, wrists, hips, knees, and ankles) in each cropped monkey image. The monkey identification network (ID Net) classified an identity (B, G, R, W, unknown, or monkey-overlap) of each cropped monkey image. We used B, G, R, and W classifications with confidence values >0.9 in the following processes. To train and test the Detection Net and Pose Net, 2,438 (training: 2,238; test: 200) images captured in the experiments, including 6,435 monkeys (training: 5,927; test: 508), were manually annotated by Cocosnet Ltd. We developed a custom annotation software capable of multi-view augmentation^13^, and reprojected 3D keypoint positions to images of all eight camera views (Movie S3). Note that we pre-trained Pose Net with the MacaquePose dataset^26^, containing 13,083 images (16,393 monkeys) captured in various naturalistic contexts. For training and test of ID Net, 16,510 (training: 16,270; test: 240) and 2,023 (training: 1,904; test: 119) cropped monkey images for Groups I and II, respectively, were manually labeled by a trained experimenter. The average precision (AP) of the trained neural networks was 0.744 for Detection Net, 0.735 for Pose Net, 0.853 for ID Net for Group I, and 0.810 for ID Net for Group II. In addition, bounding boxes were tracked in a single camera video using the ByteTrack algorithm^48^, resulting in tracklets, i.e., fragments of monkey tracks. Here, we call this type of tracklets generated with each camera view as single-view tracklets, while we call tracklets generated by integrating information from all views in the way described later as multi-view tracklets. We built this 2D video processing pipeline using popular computer vision libraries provided by OpenMMLab (openmmlab.com).

### Multi-view tracklet generation

We generated multi-view tracklets, i.e., a set of 2D monkey detections (estimated in the process described above) corresponding to each individual across views and video frames by the following processing. For cross-view matching of 2D monkey detections, we customized and used the MVPose algorithm suggested for human pose estimation^18^ (Fig 1c). To estimate optimal cross-view matches, we calculated the affinity matrix (***A***), which represents the affinity of all pairs of 2D detections in all views in a time point. We defined the affinity matrix as a weighted sum of geometric affinity (***A****^g^*) and appearance affinity (***A****^a^*), as follows:

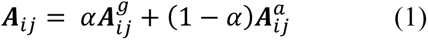

where *i* and *j* are an index of a pair of 2D monkey detections (1 ≤ *i, j* ≤ *m*; *m*, the total number of 2D monkey detections), and *α* is constant (0.8 in this study). To derive geometric affinity (***A****^g^*), we calculated the geometric distance (***D***) of a pair of 2D monkey detections as follows:

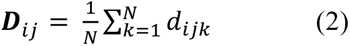

where *N* is the number of keypoints and *d_ijk_* is the distance between camera rays to the *k*-th keypoint of the *i*- and *j*-th 2D monkey detections. Only the keypoints commonly detected (prediction confidence > 0.1) in the *i*- and *j*-th 2D monkeys were included. Then, we defined geometric proximity (***C***) as follows:

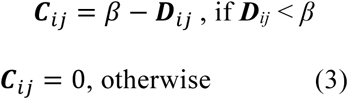

where *β* is constant (1,500 mm in this study). The geometric affinity (***A****^g^*) was obtained by mapping the proximity ***C*** to values in (0,1) with a sigmoid function. The original algorithm calculated the appearance affinity (***A****^a^*) using image feature values obtained with a person re-ID network^49^, which extracts view-invariant discriminative appearance features such as clothing and hairstyle. However, extracting such view-invariant discriminative appearance from similar-looking monkeys is difficult, so we estimated a putative ID of each single-view tracklet based on the outputs of the monkey identification network with a similar ID assignment algorithm used for the multi-view tracklets (see next section) applied separately to each view. Then, we calculated appearance affinity (***A****^a^*) as follows:

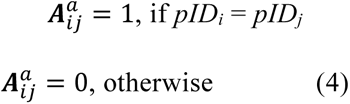

where *pID_i_*represents the putative ID of the single-view tracklet containing the *i*-th detection at the timepoint.

Finally, the permutation matrix (***P***), which represents the matching of all detection pairs, was estimated. The optimal permutation matrix should maximize the affinity and be cycle-consistent, i.e., in a matched group, there is no more than a single detection per view. According to Dong et al.^18^, such a permutation matrix was estimated by solving the following optimization problem with the alternating direction method of multipliers (ADMM):

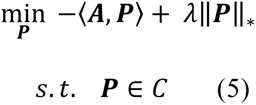

where *<**A**, **P**>* denotes the inner product of the matrices, λ represents a constant (50 in this study), and ‖***P***‖_∗_ represents the nuclear norm of ***P***. *C* represents a set of matrices satisfying the following constraints:

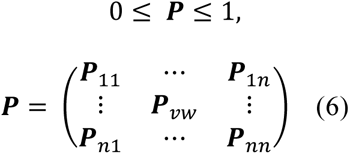

where *n* is the number of views and ***P****_vw_* is a permutation matrix that represents the matching of detection pairs of view *v* and *w*. ***P****_vw_* satisfies the following constraints:

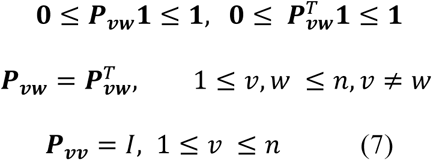

This cross-view matching was performed for one keyframe every 0.5 s to reduce computational load. Between the keyframes, cross-view and cross-frame matching was estimated using single-view tracklets and sets of the cross-view matched 2D monkey detections at keyframes. Specifically, we defined the tracking consistency of a pair of sets of the matched 2D monkey detections in neighboring keyframes as the number of single view tracklets shared by both sets (Fig 1d). Then, matching of the pairs of the sets that maximized the total tracking consistency was calculated using the Hungarian method, resulting in multi-view tracklets, i.e., a set of 2D monkey detections corresponding to an individual monkey across views and video frames.

### ID assignment to multi-view tracklets

Then, the ID of each multi-view tracklet was estimated. To compensate for occasional errors of the monkey identification network, first, we defined a detected ID of a multi-view tracklet to be valid if it passed the following criteria in a time window:

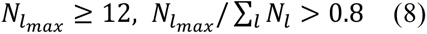

where *l* represents an index of ID (B, G, R, or W), *l_max_*represents an index of the most detected ID, and *N_l_* is the detection count of ID *l*. The valid ID of each time point was checked using a 5-s sliding window. If only a single valid ID was found in a multi-view tracklet, the ID was assigned to the tracklet. If multiple valid IDs were found, the tracklet was separated at the mid-point when different valid IDs were found. If no valid ID was found, Eq. 8 was re-examined with a time window covering the entire time range of the tracklet, and if it passed, the ID was assigned to the tracklet.

Since the linking of multi-view tracklets was interrupted when no correspondence was found between the neighboring keyframes, e.g., in the case of severe occlusion, second, we stitched the tracklets together based on their continuity using network flow optimization^15,50^. Specifically, a directed graph consisting of nodes representing tracklets and edges representing possible connections between tracklets was constructed. A pair of multi-view tracklets (nodes) separated by no more than 5 s and having at least one common single-view tracklet were connected with an edge. Each edge had a cost value equal to the distance between the monkey’s neck location at the end of one tracklet and the beginning of the other tracklet. If the assigned IDs of a tracklet pair were different, the corresponding edge was removed. If the IDs were the same, the cost value of the corresponding edge was divided by 100. In addition, sink and source nodes were assumed and connected to all tracklet nodes by the edges of a fixed amount of cost (1,000 mm in this study). Then, optimal associations between tracklets were calculated by a minimum-cost flow algorithm using the capacity_scaling() function in the NetworkX library (networkx.org), and the tracklets were stitched accordingly.

Third, the IDs were re-assigned to the stitched tracklets using the same criteria (Eq. 6). Sometimes, the ID assignment resulted in multiple tracklets having the same ID at the same time. We resolved such ID overlaps as follows: 1) if an overlapping tracklet was previously stitched and the original (before stitching) tracklet corresponding to the overlapping part had no valid ID, the tracklet was unstitched; 2) Overlapping tracklets without a non-overlapping period (Fig S3a) was excluded; and 3) Overlapping tracklets were trimmed based on the timings when the valid IDs were detected (Fig S3b). Finally, the tracklets that had no ID yet and that there were ≥3 frames in which tracklets with the other three IDs were present during the period of the tracklet were assigned to the fourth (final) ID.

### Pose filtering

We reconstructed the 3D motion of each monkey using the multi-view tracklets with the corresponding ID. For robust 3D motion estimation, we used the Anipose algorithm^19^. Briefly, 2D keypoint trajectories were Viterbi filtered in each view, and then the reasonable 3D motion of an animal was estimated by minimizing the error of body part length, change in keypoint acceleration, and reprojection error.

### Detection of social behavioral events

We calculated basic behavioral parameters using the estimated 3D motions of monkeys to detect behavioral events. Specifically, we determined the position (neck; mid-point between left and right shoulder), speed, and face direction (vector from the mid-point of the left and right ears to the nose, rotated upward by 35°) of each monkey. We also calculated the distance and approaching/leaving speed (speed along the axis connecting the pair) for each pair of monkeys. We lowpass filtered these parameters with a cut-off frequency of 0.5 Hz. In addition, we classified body-centered postures in a feature space obtained with principal component analysis to detect sitting, lying (Fig S4a), and typical forelimb posture associated with grooming (Fig S4b). The social behavioral events were detected (Fig 3a) with these parameters according to the definitions shown in Text S1.

### Performance validation

To examine the performance of 3D pose estimation, we annotated the 3D poses of monkeys with one frame every 12 s from eight 5-min video clips that did not overlap with the training dataset. The error of each keypoint between a manually annotated and automatically estimated pose was calculated when the corresponding instance was found (root mean square error of all keypoints < 500 mm). To assess the performance of 3D tracking, we annotated the 3D neck positions of monkeys with IDs in 2.5 frames/s from the same video clips. Manually annotated instances and automatically estimated instances in each frame were matched by the Hungarian method to minimize the total distance between the necks of the matched pairs. If the distance of a matched pair was <400 mm and the IDs of the pair were coincident, it was counted as successful (true positive [TP]). The number of false positives (FP) and false negatives (FN) was calculated by subtracting TP from the total number of automatically estimated or manually annotated instances, respectively.

We tested the performance of social behavioral event detection using another dataset. Specifically, two human experts annotated the occurrence and direction (from which monkey to which monkey) of Chase, Groom, and Mount between a given pair in 10-s video clips, and annotations common to both experts were used as the ground truth. TP, FP, and FN cases were counted by comparing the annotated results with the automatic estimation. Seven hundred and one video clips (Dataset A) were selected based on successful tracking (both monkeys were tracked for >50% of the time of the clip) and potential of interaction (the distance between the monkeys became <1 m at least once in the clip). Since Mount was a rare event, we added 11 video clips (Dataset B) when Mount was detected automatically. Both datasets were shuffled and presented to human experts for annotation. In the performance evaluation, we used Dataset A for Chase and Groom and Dataset B for Mount.

We defined and calculated Precision and Recall values as TP / (TP + FP) and TP / (TP + FN), respectively, to evaluate the performance of ID tracking and behavioral event detection.

### Behavioral data analysis

The behavioral data analyses were mainly focused on 08:30–17:30 on each group’s last 8 recording days. We analyzed a consecutive 5-min session every 30 min (08:30–08:35, 09:00– 09:05…, 17:00–17:05) to reduce computational load. We excluded a session from the analysis if an experimenter was inside or in front of the cage. Only sessions with >30 s detection of all four monkeys were used to analyze their interactions. As a result, we obtained 58 and 111 sessions for Groups I and II, respectively (Table S2).

We used an SVM to predict monkey IDs from behavioral patterns to assess individual differences in the behavioral patterns. Twenty sessions (100 min) were combined and used as input to the SVM, and Leave-one-out cross-validation was used to evaluate the performance of the SVM. Specifically, we used a set of 20 randomly sampled sessions as test data. From the rest of the data, we randomly sampled 100 combinations of 20 sessions and used them for training the SVM. Then, we compared the trained SVM’s ID predictions on the test data with the correct IDs. This training-test process was repeated 100 times, and the accuracy of ID prediction by the SVM was calculated.

Statistical tests were performed using SPSS and MATLAB. The significance threshold was set to 0.05.

## Supporting information

Movie S1: An example of 3D motion capture

Movie S2: Examples of Chase, Groom, and Mount events

Movie S3: Example use of the custom 3D annotation software

## Data availability

The training datasets will be made available in a public repository at the time of publication.

## Code availability

The original codes will be made available in a public repository at the time of publication.

## Acknowledgments

We are grateful to A. Sharif, A. Badarch, and E. Sumiya for their technical assistance. This work was supported by MEXT/JSPS KAKENHI Grant Numbers 16H06534 and 22H05157 to J.M., 22K07325 to T.K., 16H06534 to T.Sh., 19H05467 to M.T., 22K19480, 22H05157, and 23H02781 to K.I., by JST Grant Number JPMJMS2295-12 to K.I., by the Takeda Science Foundation to J.M., T.Se., H.Nm., and H.Nj., and by the National Institutes of Natural Sciences (NINS) Joint Research Grant Number 01111901 to J.M., Y.G., T.Sh., and K.I.

## Author contributions

J.M., T.K., Y.G., T.Sh., and K.I. designed the experiments. J.M., T.K., K.K., J.G., N.S., M.M., and K.I. designed and constructed the recording setup. T.K., K.K., N.S., A.K., and M.M. recorded the data. J.M., N.S., M.M., and K.I. created the annotated dataset. J.M., T.K., S.B.N., and K.I. constructed the motion capture pipeline. J.M., T.K., T.Se., and K.I. analyzed the data. J.M., T.K., H.Nm., T.Se., H.Nj., M.T., and K.I. wrote the manuscript. All authors discussed the data and commented on the manuscript.

## Competing interests

The authors declare no competing interests.

## Figures

**Figure S1:**
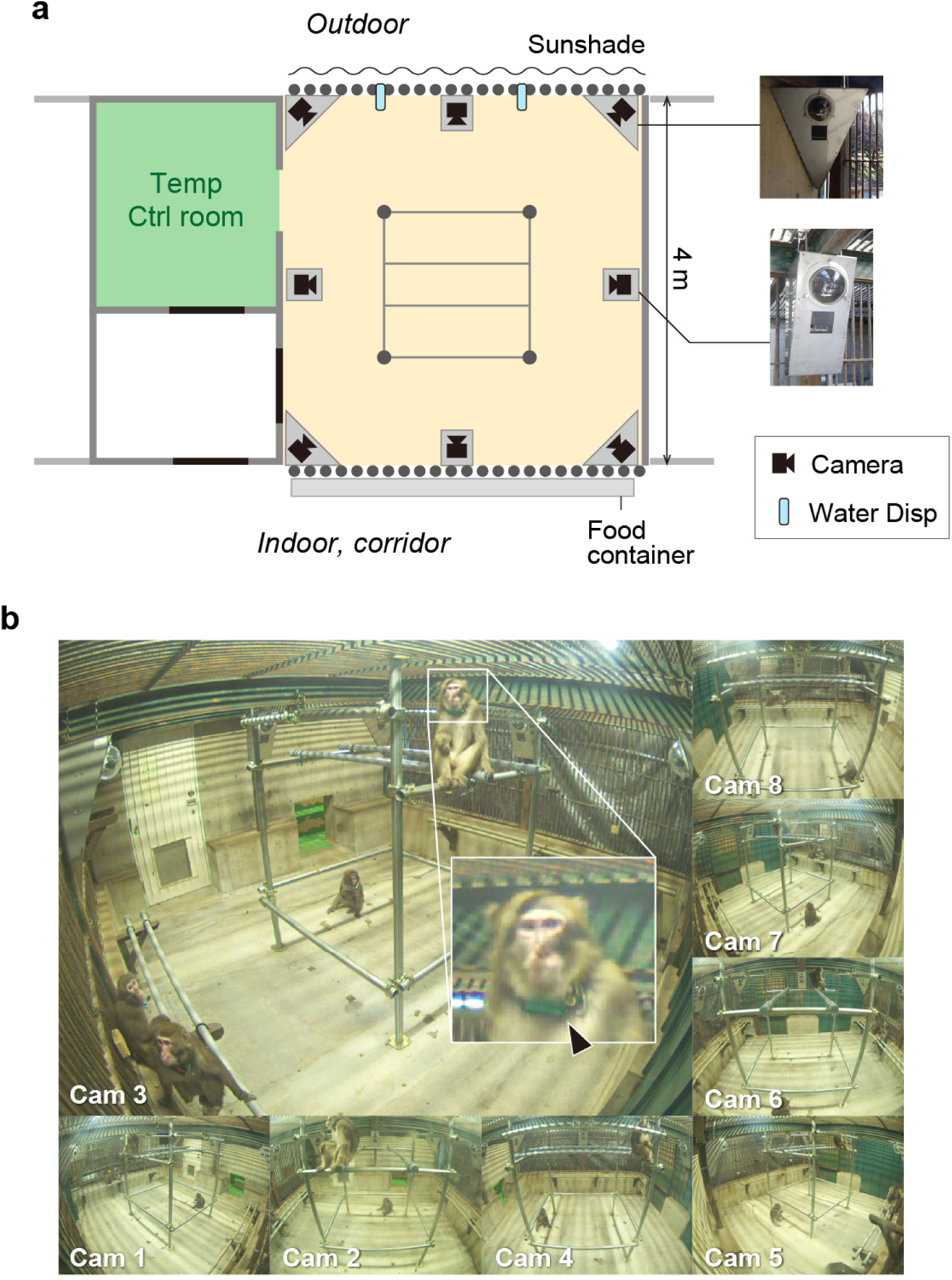
Recording setup. (**a**) Schema of the recording setup. The square in the center of the cage represents a jungle gym. Temp Ctrl room, temperature-controlled room. Water Disp, water dispenser. Inset pictures show the camera housings. (**b**) An example set of images captured simultaneously from the eight cameras. The inset shows the color tag (arrowhead) for monkey identification.

**Figure S2:**
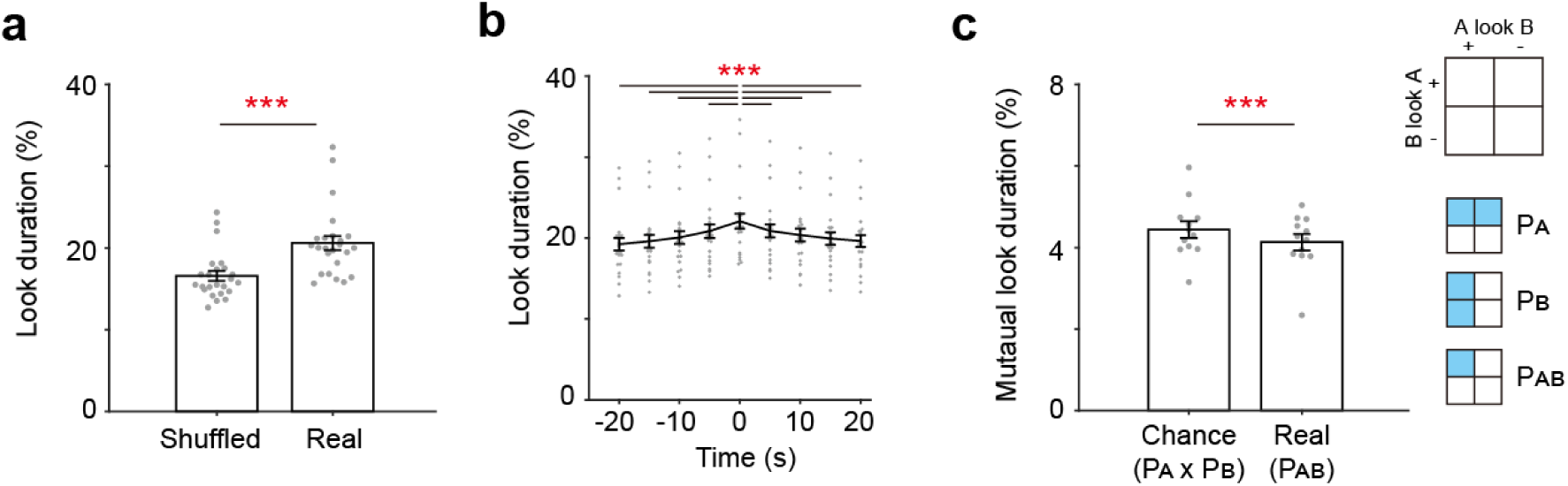
Looking pattern is not random. (**a**) Comparison of the mean Look duration with (Shuffled) and without (Real) shuffling of the subjects’ motion data across recording sessions. \*\*\**p* < 0.001, paired *t*-test. *n* = 24 pairs. (**b**) Mean Look duration with temporally shifting subjects’ motion data. Horizontal axis, the amount of time shift. \*\*\**p* < 0.001, paired *t*-test compared with the value at time = 0 with Bonferroni’s correction. *n* = 24 pairs. (**c**) Comparison of the actual mutual look duration (Real) with the chance-level duration (Chance). The chance level was calculated by multiplying the probability of one monkey A looking at another monkey B (corresponding to P_A_ in inset) and *vice versa* (P_B_). \*\*\**p* < 0.001, paired *t*-test. *n* = 12 pairs.

**Figure S3:**
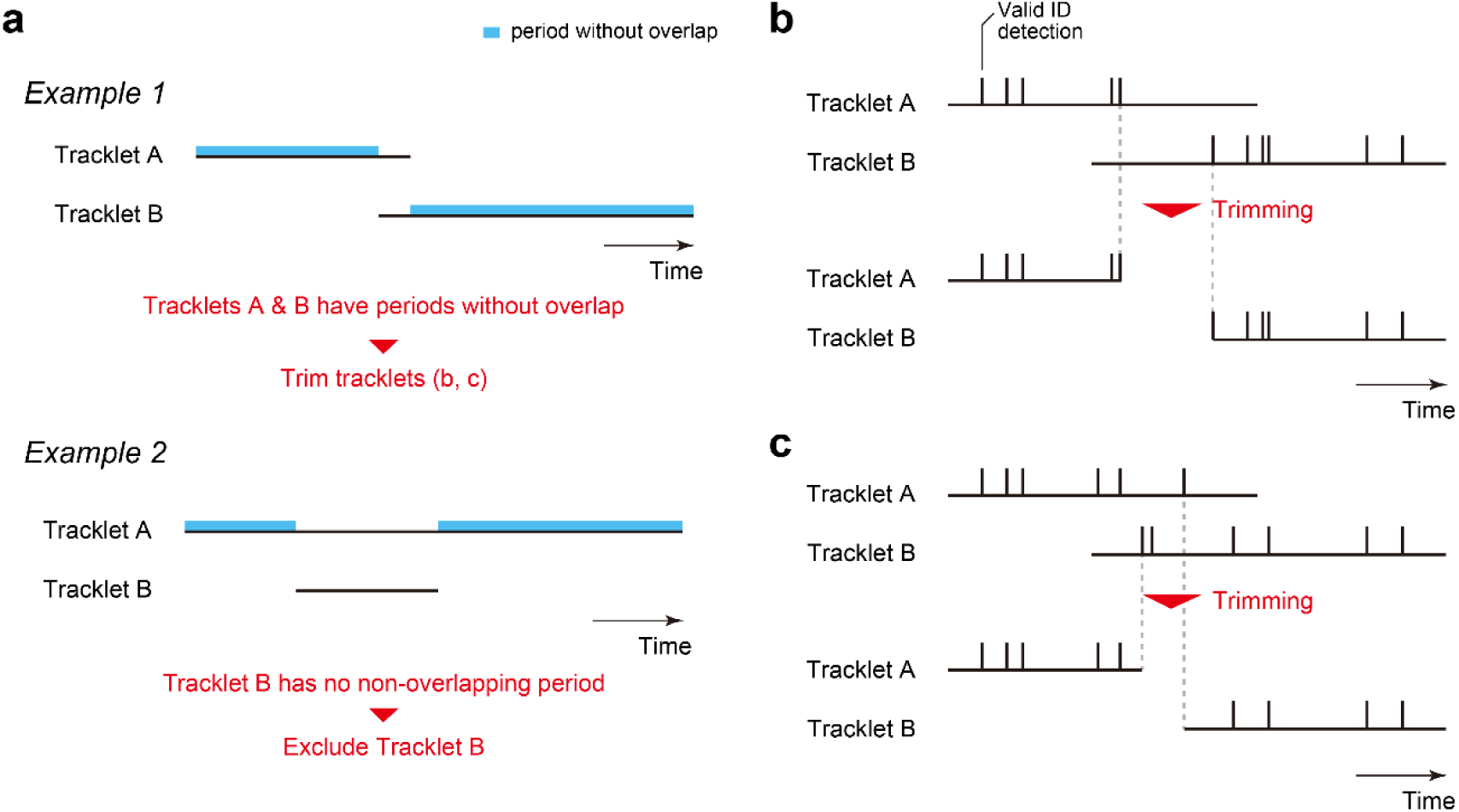
Resolving temporal overlap of tracklets with the same ID. (**a**) Overlapping tracklets in which both tracks have a non-overlapping period (light blue area) are trimmed as described in **b** and **c** (Example 1). A tracklet without a non-overlapping period (Tracklet B in Example 2) was excluded. (**b** and **c**) Tracklet trimming. If the last valid ID detection of the preceding tracklet was before the first valid ID detection of the following tracklet, the preceding and following tracklets are trimmed to their own last and first detection time, respectively (b). Otherwise, the preceding and following tracklets are trimmed at the other’s first and last detection time, respectively (c).

**Figure S4:**
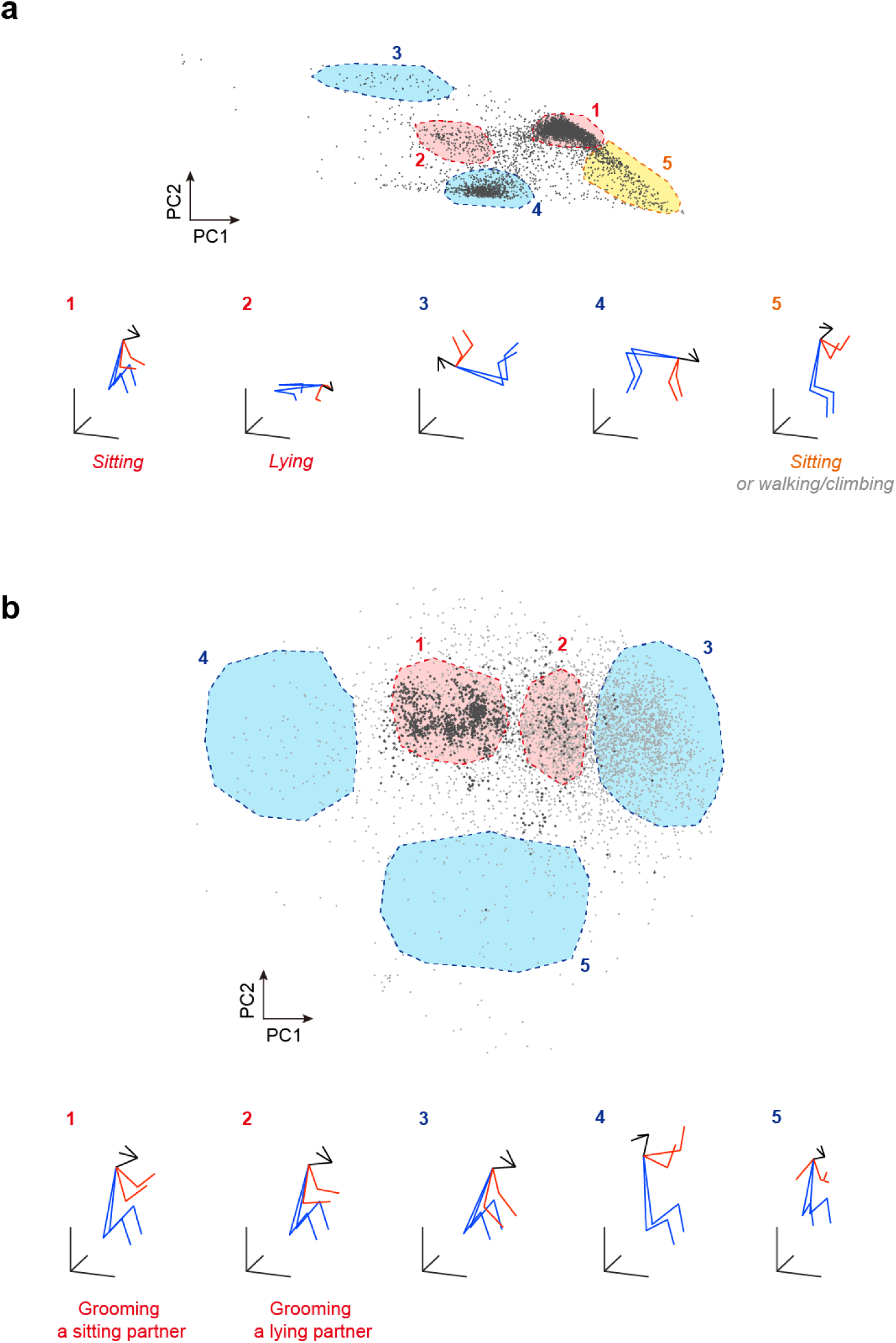
Classification of postures using principal component analysis (PCA). (**a**) To classify sitting and lying postures, the vertical positions of keypoints of the lower body (left and right hips, knees, and ankles) are reduced to two dimensions by PCA (top). The boundaries of clusters in the feature space corresponding to sitting (1, 5) and lying (2) are drawn for the classification. Since the postures in cluster 5 appear during walking on two legs and climbing a wall, as well as sitting with the legs down, the posture in cluster 5 was classified as sitting only when the hip center is located at a place where a monkey can sit (within 100 mm where the hip center of the postures in cluster 1 are observed). Bottom, visualization of average posture in each cluster. (**b**) To classify postures associated with grooming, 3D positions of keypoints of the upper body (nose, left and right ears, elbows, and wrists; shoulders were not considered since they were fixed in the body-centered coordinate) for monkeys in Proximity are reduced to two dimensions by PCA (top). Some grooming behaviors were manually annotated by visual inspection of videos, and the corresponding postures are plotted with dark dots. From the distribution of the dark plots, the two clusters (1, 2), corresponding to the postures during grooming for a sitting and lying partner, were identified and used for classification. Bottom, visualization of the average posture of the clusters (1, 2) and those of the other areas in the feature space (3–5).

## Tables

**Table S1:**
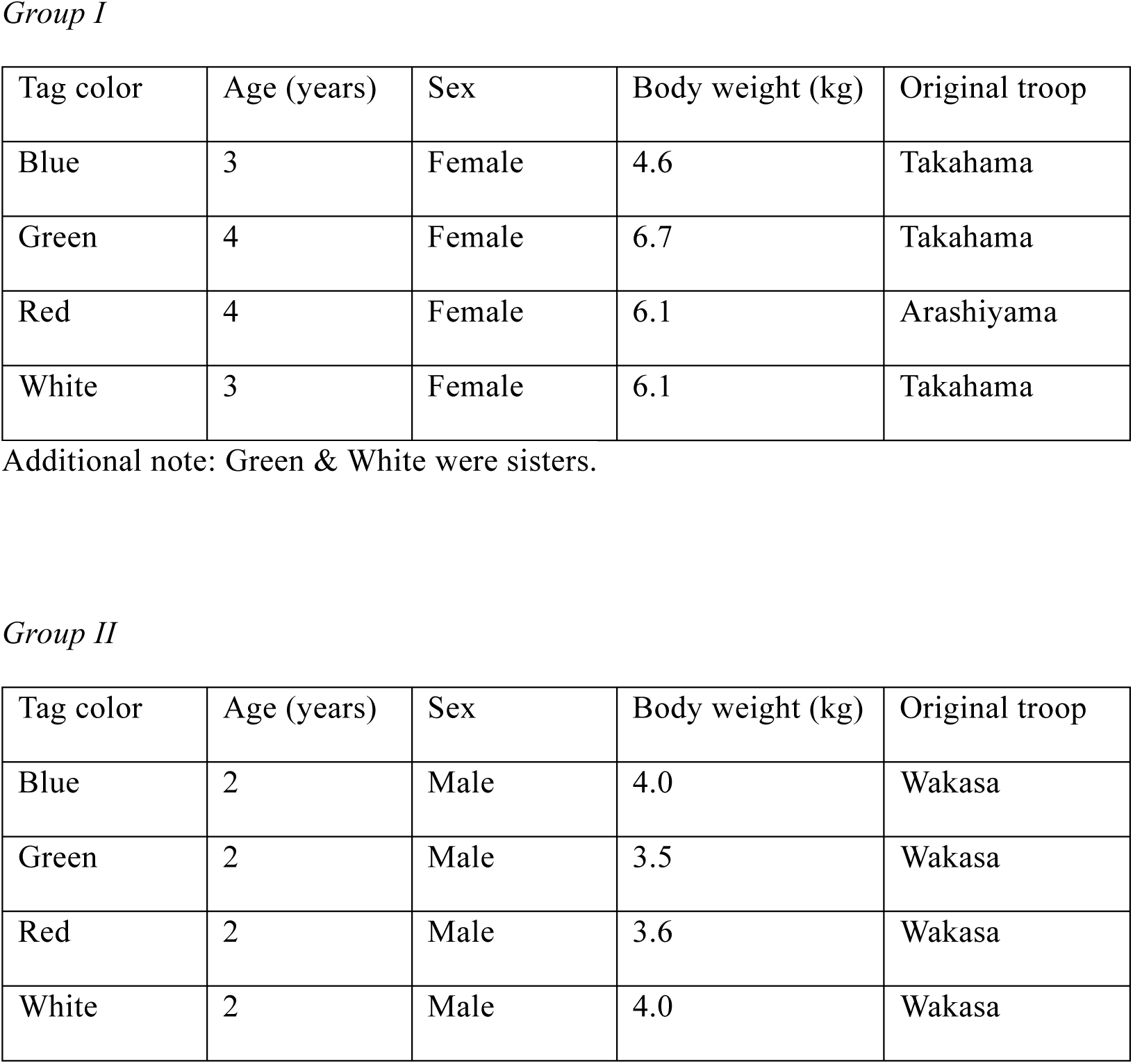
Monkey profiles.

*Group I*
| Tag color | Age (years) | Sex | Body weight (kg) | Original troop |
| --- | --- | --- | --- | --- |
| Blue | 3 | Female | 4.6 | Takahama |
| Green | 4 | Female | 6.7 | Takahama |
| Red | 4 | Female | 6.1 | Arashiyama |
| White | 3 | Female | 6.1 | Takahama |

*Group II*
| Tag color | Age (years) | Sex | Body weight (kg) | Original troop |
| --- | --- | --- | --- | --- |
| Blue | 2 | Male | 4.0 | Wakasa |
| Green | 2 | Male | 3.5 | Wakasa |
| Red | 2 | Male | 3.6 | Wakasa |
| White | 2 | Male | 4.0 | Wakasa |

**Table S2:**
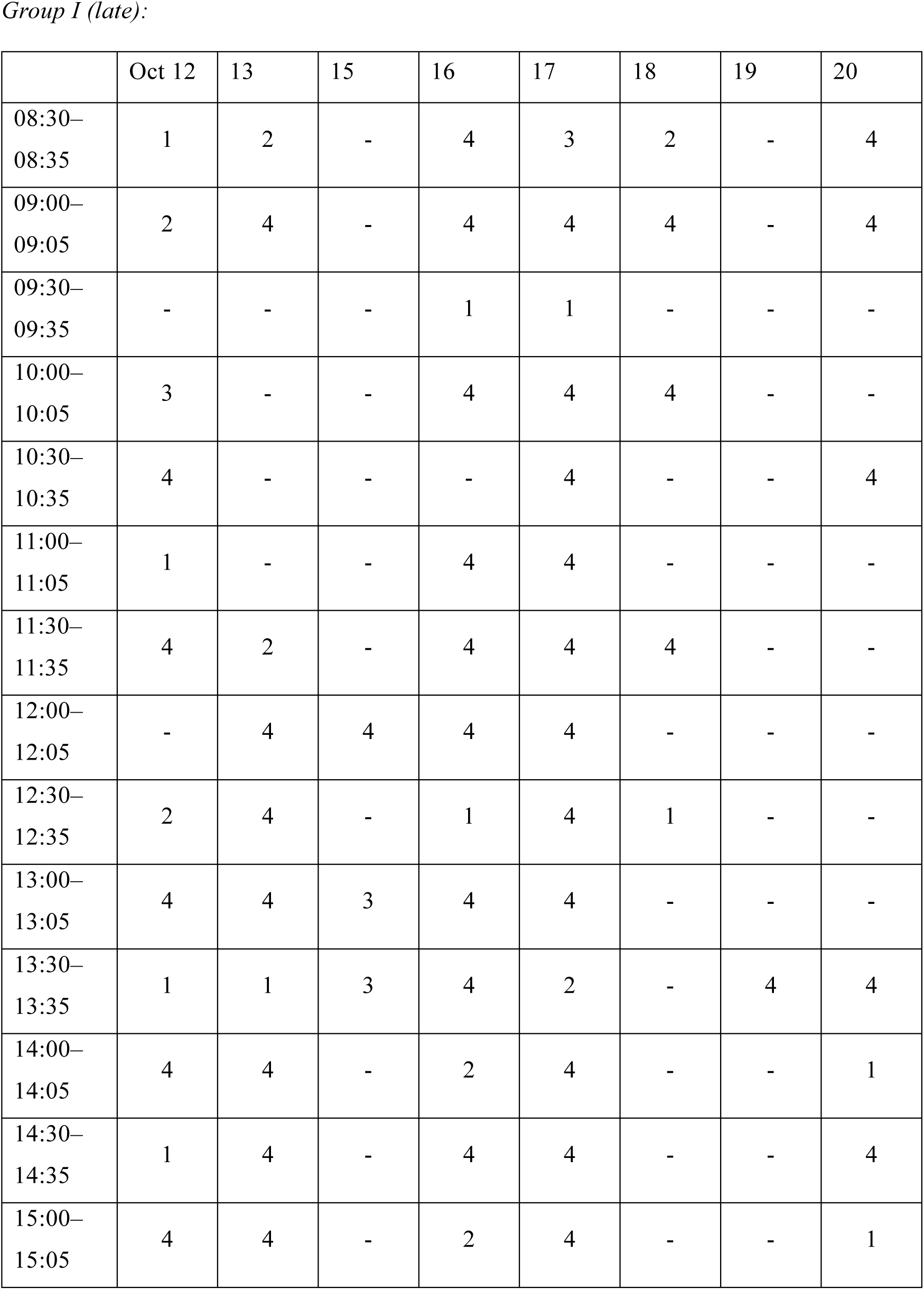
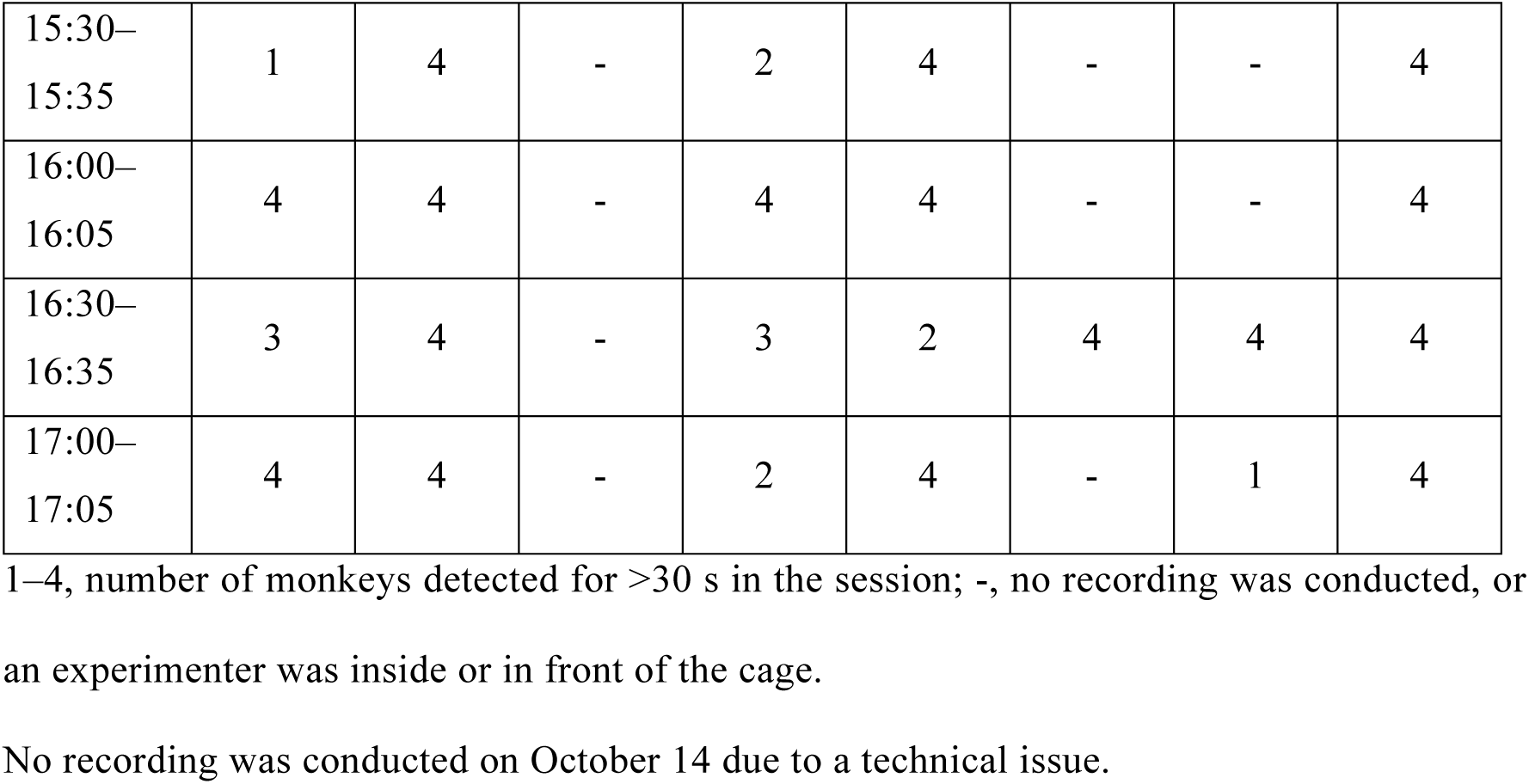
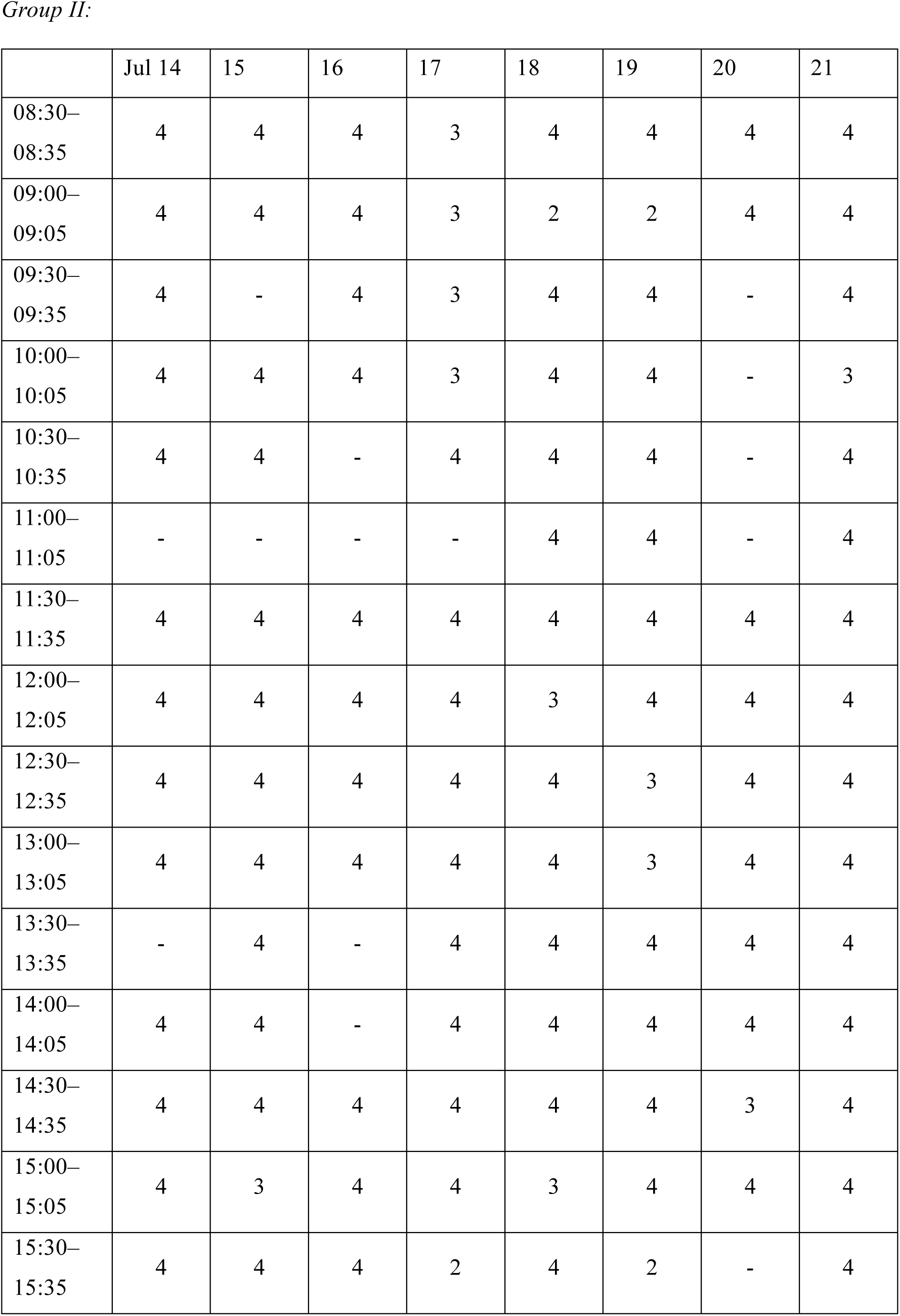
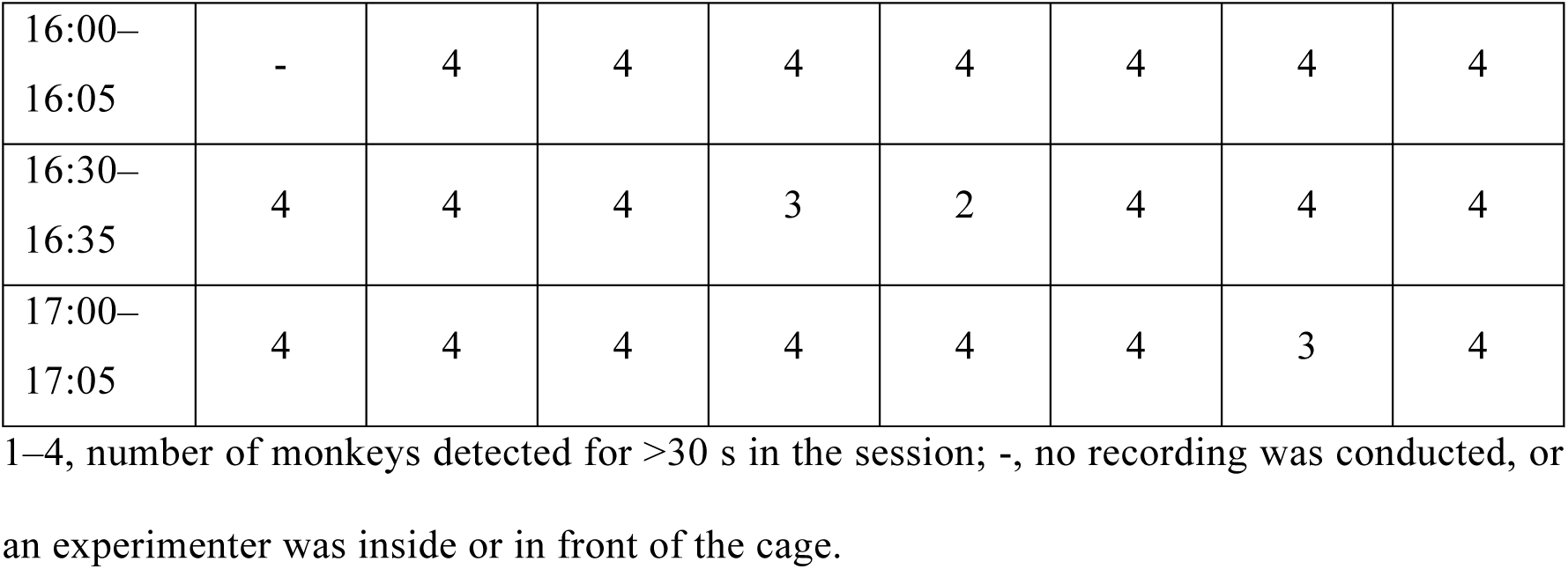
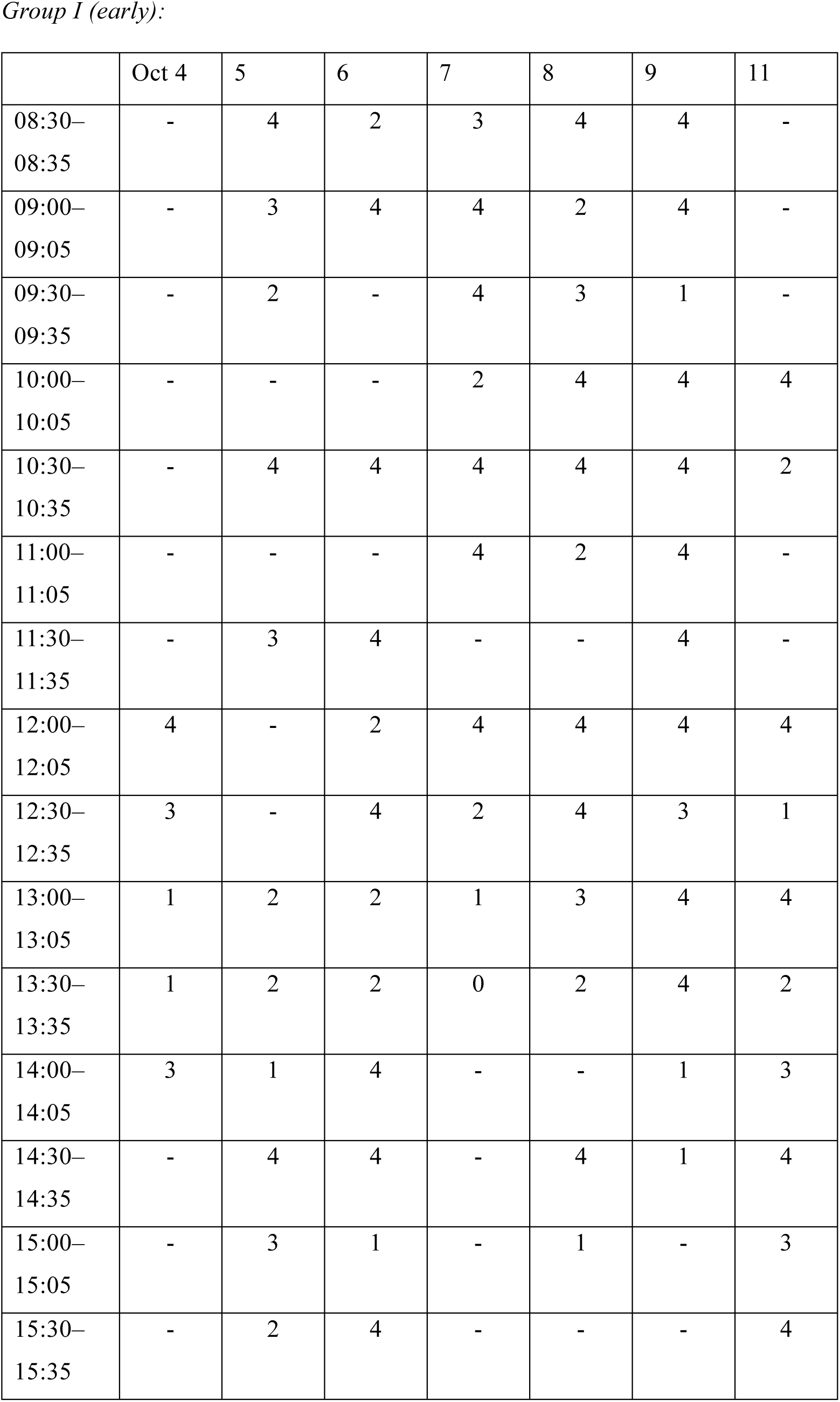
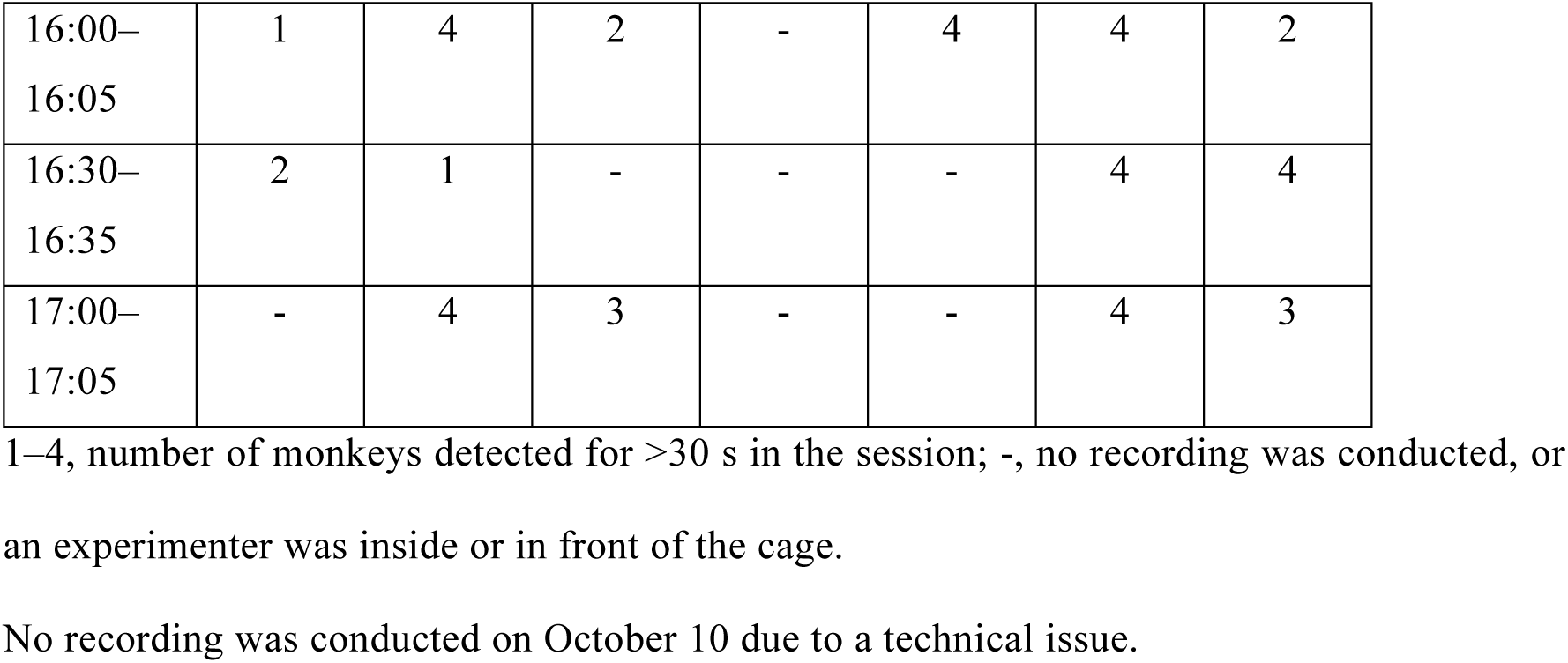
Recording sessions.

*Group I (late):*
|  | Oct 12 | 13 | 15 | 16 | 17 | 18 | 19 | 20 |
| --- | --- | --- | --- | --- | --- | --- | --- | --- |
| 08:30–<br>08:35 | 1 | 2 | - | 4 | 3 | 2 | - | 4 |
| 09:00–<br>09:05 | 2 | 4 | - | 4 | 4 | 4 | - | 4 |
| 09:30–<br>09:35 | - | - | - | 1 | 1 | - | - | - |
| 10:00–<br>10:05 | 3 | - | - | 4 | 4 | 4 | - | - |
| 10:30–<br>10:35 | 4 | - | - | - | 4 | - | - | 4 |
| 11:00–<br>11:05 | 1 | - | - | 4 | 4 | - | - | - |
| 11:30–<br>11:35 | 4 | 2 | - | 4 | 4 | 4 | - | - |
| 12:00–<br>12:05 | - | 4 | 4 | 4 | 4 | - | - | - |
| 12:30–<br>12:35 | 2 | 4 | - | 1 | 4 | 1 | - | - |
| 13:00–<br>13:05 | 4 | 4 | 3 | 4 | 4 | - | - | - |
| 13:30–<br>13:35 | 1 | 1 | 3 | 4 | 2 | - | 4 | 4 |
| 14:00–<br>14:05 | 4 | 4 | - | 2 | 4 | - | - | 1 |
| 14:30–<br>14:35 | 1 | 4 | - | 4 | 4 | - | - | 4 |
| 15:00–<br>15:05 | 4 | 4 | - | 2 | 4 | - | - | 1 |
| 15:30–<br>15:35 | 1 | 4 | - | 2 | 4 | - | - | 4 |
| 16:00–<br>16:05 | 4 | 4 | - | 4 | 4 | - | - | 4 |
| 16:30–<br>16:35 | 3 | 4 | - | 3 | 2 | 4 | 4 | 4 |
| 17:00–<br>17:05 | 4 | 4 | - | 2 | 4 | - | 1 | 4 |

|  | Jul 14 | 15 | 16 | 17 | 18 | 19 | 20 | 21 |
| --- | --- | --- | --- | --- | --- | --- | --- | --- |
| 08:30–<br>08:35 | 4 | 4 | 4 | 3 | 4 | 4 | 4 | 4 |
| 09:00–<br>09:05 | 4 | 4 | 4 | 3 | 2 | 2 | 4 | 4 |
| 09:30–<br>09:35 | 4 | - | 4 | 3 | 4 | 4 | - | 4 |
| 10:00–<br>10:05 | 4 | 4 | 4 | 3 | 4 | 4 | - | 3 |
| 10:30–<br>10:35 | 4 | 4 | - | 4 | 4 | 4 | - | 4 |
| 11:00–<br>11:05 | - | - | - | - | 4 | 4 | - | 4 |
| 11:30–<br>11:35 | 4 | 4 | 4 | 4 | 4 | 4 | 4 | 4 |
| 12:00–<br>12:05 | 4 | 4 | 4 | 4 | 3 | 4 | 4 | 4 |
| 12:30–<br>12:35 | 4 | 4 | 4 | 4 | 4 | 3 | 4 | 4 |
| 13:00–<br>13:05 | 4 | 4 | 4 | 4 | 4 | 3 | 4 | 4 |
| 13:30–<br>13:35 | - | 4 | - | 4 | 4 | 4 | 4 | 4 |
| 14:00–<br>14:05 | 4 | 4 | - | 4 | 4 | 4 | 4 | 4 |
| 14:30–<br>14:35 | 4 | 4 | 4 | 4 | 4 | 4 | 3 | 4 |
| 15:00–<br>15:05 | 4 | 3 | 4 | 4 | 3 | 4 | 4 | 4 |
| 15:30–<br>15:35 | 4 | 4 | 4 | 2 | 4 | 2 | - | 4 |
| 16:00–<br>16:05 | - | 4 | 4 | 4 | 4 | 4 | 4 | 4 |
| 16:30–<br>16:35 | 4 | 4 | 4 | 3 | 2 | 4 | 4 | 4 |
| 17:00–<br>17:05 | 4 | 4 | 4 | 4 | 4 | 4 | 3 | 4 |

|  | Oct 4 | 5 | 6 | 7 | 8 | 9 | 11 |
| --- | --- | --- | --- | --- | --- | --- | --- |
| 08:30–<br>08:35 | - | 4 | 2 | 3 | 4 | 4 | - |
| 09:00–<br>09:05 | - | 3 | 4 | 4 | 2 | 4 | - |
| 09:30–<br>09:35 | - | 2 | - | 4 | 3 | 1 | - |
| 10:00–<br>10:05 | - | - | - | 2 | 4 | 4 | 4 |
| 10:30–<br>10:35 | - | 4 | 4 | 4 | 4 | 4 | 2 |
| 11:00–<br>11:05 | - | - | - | 4 | 2 | 4 | - |
| 11:30–<br>11:35 | - | 3 | 4 | - | - | 4 | - |
| 12:00–<br>12:05 | 4 | - | 2 | 4 | 4 | 4 | 4 |
| 12:30–<br>12:35 | 3 | - | 4 | 2 | 4 | 3 | 1 |
| 13:00–<br>13:05 | 1 | 2 | 2 | 1 | 3 | 4 | 4 |
| 13:30–<br>13:35 | 1 | 2 | 2 | 0 | 2 | 4 | 2 |
| 14:00–<br>14:05 | 3 | 1 | 4 | - | - | 1 | 3 |
| 14:30–<br>14:35 | - | 4 | 4 | - | 4 | 1 | 4 |
| 15:00–<br>15:05 | - | 3 | 1 | - | 1 | - | 3 |
| 15:30–<br>15:35 | - | 2 | 4 | - | - | - | 4 |
| 16:00–<br>16:05 | 1 | 4 | 2 | - | 4 | 4 | 2 |
| 16:30–<br>16:35 | 2 | 1 | - | - | - | 4 | 4 |
| 17:00–<br>17:05 | - | 4 | 3 | - | - | 4 | 3 |

1–4, number of monkeys detected for >30 s in the session; -, no recording was conducted, or
an experimenter was inside or in front of the cage.
No recording was conducted on October 10 due to a technical issue.

**Table S3:**
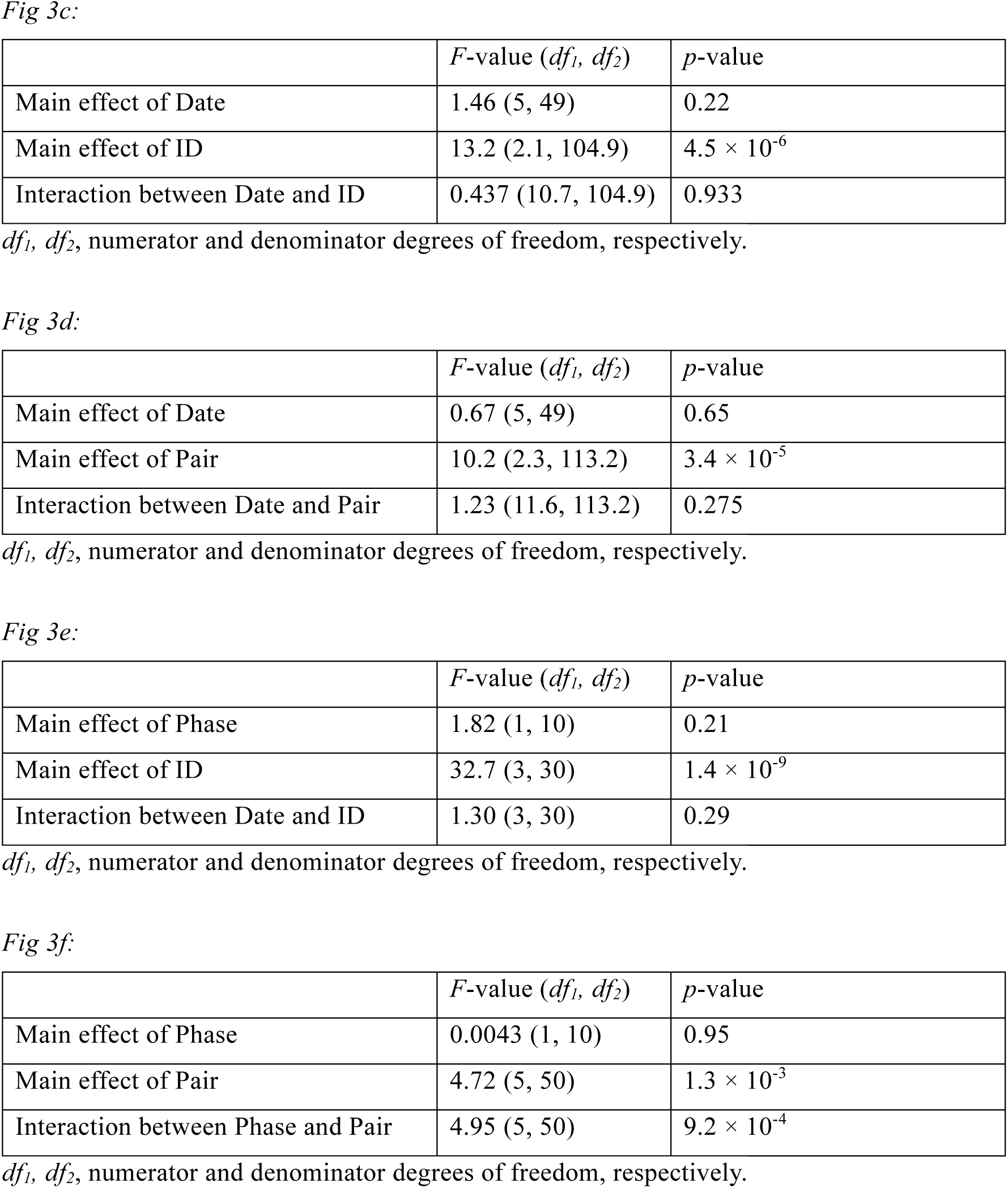
Analysis of variance results for Fig 3.

|  | *F*-value (*df*<sub>1</sub>, *df*<sub>2</sub>) | *p*-value |
| --- | --- | --- |
| Main effect of Date | 1.46 (5, 49) | 0.22 |
| Main effect of ID | 13.2 (2.1, 104.9) | $4.5 \times 10^{-6}$ |
| Interaction between Date and ID | 0.437 (10.7, 104.9) | 0.933 |

|  | *F*-value (*df*<sub>1</sub>, *df*<sub>2</sub>) | *p*-value |
| --- | --- | --- |
| Main effect of Date | 0.67 (5, 49) | 0.65 |
| Main effect of Pair | 10.2 (2.3, 113.2) | $3.4 \times 10^{-5}$ |
| Interaction between Date and Pair | 1.23 (11.6, 113.2) | 0.275 |

|  | *F*-value (*df*<sub>1</sub>, *df*<sub>2</sub>) | *p*-value |
| --- | --- | --- |
| Main effect of Phase | 1.82 (1, 10) | 0.21 |
| Main effect of ID | 32.7 (3, 30) | $1.4 \times 10^{-9}$ |
| Interaction between Date and ID | 1.30 (3, 30) | 0.29 |

|  | *F*-value (*df*<sub>1</sub>, *df*<sub>2</sub>) | *p*-value |
| --- | --- | --- |
| Main effect of Phase | 0.0043 (1, 10) | 0.95 |
| Main effect of Pair | 4.72 (5, 50) | $1.3 \times 10^{-3}$ |
| Interaction between Phase and Pair | 4.95 (5, 50) | $9.2 \times 10^{-4}$ |

*df*<sub>1</sub>, *df*<sub>2</sub>, numerator and denominator degrees of freedom, respectively.

**Table S4:** Filter parameters for behavioral events.

| | $I_{max}$ | $D_{min}$ |
| --- | --- | --- |
| *Proximity* | 0.5 | 0.5 |
| *Grooming* | 1.0 | 1.0 |
| *Mount* | 1.5 | 0.17 |
| *Chase* | 0.5 | 0.25 |
| *Glare* | 1.0 | 1.0 |
| *Grab/Push* | 2.0 | 0.08 |
| *Pounce* | 0.5 | 0.17 |

unit: seconds

## Movies

**Movie S1: An example of 3D motion capture.**

**Movie S2: Examples of Chase, Groom, and Mount events.**

**Movie S3: Example use of the custom 3D annotation software.**

## Supplementary text

### Text S1: Definitions of behavioral events

Below, for directional behavioral events, we call the subject of the behavior as Monkey X and the object as Monkey Y.

***Look*:** The angle between the face direction of Monkey X and a vector from the nose of Monkey X to the neck of Monkey Y is <40°.

***Proximity***: The monkey pair show a sitting or lying posture (Fig S4a) and do not move fast (speed < 1,500 mm/s). The distance between the monkeys is <600 mm.

***Grooming***: Monkey Y is either sitting or lying. Monkey X is in one of the grooming-associated postures (Fig S4b) corresponding to the posture of Monkey Y. The distance between the monkeys is <500 mm, and neither monkey moves fast (speed < 1,500 mm/s). The angle between the face direction of Monkey X and a vector from the nose of Monkey X to the neck of Monkey Y is <70°, i.e., Monkey Y is in front of Monkey X.

***Mount***: The distance between the hips of the monkeys is <300 mm. The relative neck height of Monkey X from that of Monkey Y is >200 mm. The relative wrist height of Monkey X from the neck of Monkey Y is <200 mm. The mean distance between the wrists of Monkey X and the neck of Monkey Y is <400 mm. The relative ankle height of Monkey Y from the neck of Monkey Y is <-150 mm. Neither monkey is in a sitting or lying posture.

***Chase***: Monkey X approaches Monkey Y (approaching speed > 1,000 mm/s), while Monkey Y leaves from Monkey X (leaving speed > 1,000 mm/s). Both monkeys move fast (speed > 1,500 mm/s). The distance between the monkeys is <2,500 mm.

***Glare***: The monkeys *Look* at each other. The distance between the monkeys is <1,500 mm. Neither monkey moves fast (speed < 1,500 mm/s). At least one of the monkeys is not sitting or lying.

***Grab/Push***: The distance from either of Monkey X’s wrists to the neck of Monkey Y is <200 mm. The distance between the monkeys is >350 mm. The posture of Monkey X is not sitting or lying. Monkey X is not *Mounting* Monkey Y.

***Pounce***: The distance between the monkeys is <600 mm. Neither monkey moves fast (speed < 1,500 mm/s). At least one of the monkeys is not sitting or lying. The approach speed from Monkey X to Monkey Y is >0 and <1,500 mm/s. The acceleration of the approach speed is >2,500 mm/s.

Since the monkeys were smaller in Group II than in Group I, the distance, speed, and acceleration thresholds above were normalized by multiplying the values by 0.82 for Group II. After the detection of each event in each frame, except for *Look*, the event markers were filtered according to the following rules: if the same type of event occurred within an interval of less than *I_max_*, the event was considered to have continued during the interval; and if the duration of an event was less than *D_min_*, the event was ignored. The filter parameters for each event are shown in Table S4.

## Notes

### Competing Interest Statement

The authors have declared no competing interest.

